# MYC Suppresses Mitochondrial Function in Hepatocellular Carcinoma through NRF1-Driven Receptor-Mediated Mitophagy

**DOI:** 10.64898/2026.07.13.738248

**Authors:** Nicholas P. Lesner, Laura C. Kim, Spencer D. Shelton, Madelyn Landis, Xuanyan Cai, Denise Zheng, Tanay Parnaik, Caroline Bartman, Brian Keith, M. Celeste Simon

**Affiliations:** Abramson Family Cancer Research Institute, Perelman School of Medicine, University of Pennsylvania, Philadelphia, PA. 19104; Department of Cell and Developmental Biology, University of Pennsylvania, Philadelphia, PA. 19104; Division of Hematology/Oncology, Boston Children’s Hospital, Harvard Medical School, Boston, MA 02115, USA; Department of Pediatric Oncology, Dana-Farber Cancer Institute, Harvard Medical School, Boston, MA 02215, USA; Department of Cancer Biology, University of Pennsylvania, Philadelphia, PA. 19104; Systems Pharmacology and Translational Therapeutics, University of Pennsylvania, Philadelphia, PA. 19104

**Keywords:** MYC, Mitophagy, Hepatocellular Carcinoma, NRF1, Oxidative Phosphorylation

## Abstract

MYC is a canonical driver of mitochondrial biogenesis, yet its role in mitochondrial homeostasis in cancer remains incompletely understood. Here we show that MYC paradoxically suppresses mitochondrial function in hepatocellular carcinoma (HCC) through elevated receptor-mediated mitophagy. Pan-cancer TCGA analysis revealed that MYC activity exhibited discordant nuclear and mtDNA-encoded mitochondrial gene expression, particularly in liver cancer. Using genetically defined murine HCC models, we demonstrate that MYC, but not β-catenin (*CTNNB1*), drives mitochondrial fragmentation and turnover through a *MYC-NRF1-FUNDC1/PGAM5/DNM1L* (DRP1) transcriptional axis. Genetic or pharmacologic inhibition of this axis reduced mitophagy and restored oxidative phosphorylation in tumors while extending survival of tumor-bearing mice. Tumors expressing yeast NDI1, which bypasses Complex I to promote electron transport, restored OXPHOS and reduced tumor burden independent of mitophagy. Our data demonstrate that suppression of mitochondrial OXPHOS unexpectedly supports HCC tumor growth and implicates mitochondrial quality control as a novel therapeutic vulnerability in patients with *MYC*-amplified HCC.

**Significance:** Contrary to its canonical role in mitochondrial biogenesis, MYC suppresses mitochondrial function in liver cancer through a defined NRF1-driven mitophagy program. Targeting this pathway genetically or pharmacologically extends survival in tumor-bearing mice. These findings identify mitochondrial quality control as a therapeutic vulnerability in *MYC*-amplified hepatocellular carcinoma patients.

## Introduction

Incidence and mortality of liver cancer continue to rise across the globe.^1^ Hepatocellular carcinomas (HCCs), the most common form, are genetically heterogeneous with no approved therapies targeting oncogenic drivers.^2^ Treatment options remain limited, and patients are prone to developing drug resistance, leading to disease relapse.^3^ Murine models accurately reflecting the heterogeneity observed in HCC patients may identify novel liver cancer vulnerabilities and result in effective new treatments. Two commonly altered HCC oncogenes are c-MYC (*MYC*, amplification) and β-catenin (*CTNNB1,* mutation), which produce tumors with distinct biochemical and clinical features.^4,5^

Both MYC and β-catenin are transcription factors that upregulate numerous target genes, including those encoding metabolic enzymes critical to mitochondrial function.^6,7^ MYC is canonically associated with mitochondrial biogenesis, yet its function in the hypoxic, metabolically dysregulated HCC microenvironment is incompletely understood. The mitochondrion is a critical metabolic hub whose dysfunction can trigger apoptosis and immune activation through ROS-mediated damage. Cells counter this through mitochondrial quality control, encompassing biogenesis, dynamics (fission and fusion), and mitophagy. Mitochondrial fusion is mediated primarily by three large GTPases, mitofusin 1 and 2 (MFN1 and MFN2) and optic atrophy 1 (OPA1), along with other accessory factors.^8^ Conversely, mitochondrial fission is regulated by phosphorylation of Dynamin-related Protein 1 (DRP1, encoded by the *DNM1L* gene) along with other receptor proteins (**Figure 1A**). Fission is necessary for efficient mitochondrial clearance via autophagosomes, as enlarged mitochondria are refractory to mitophagy.^9,10^

**Figure 1.**
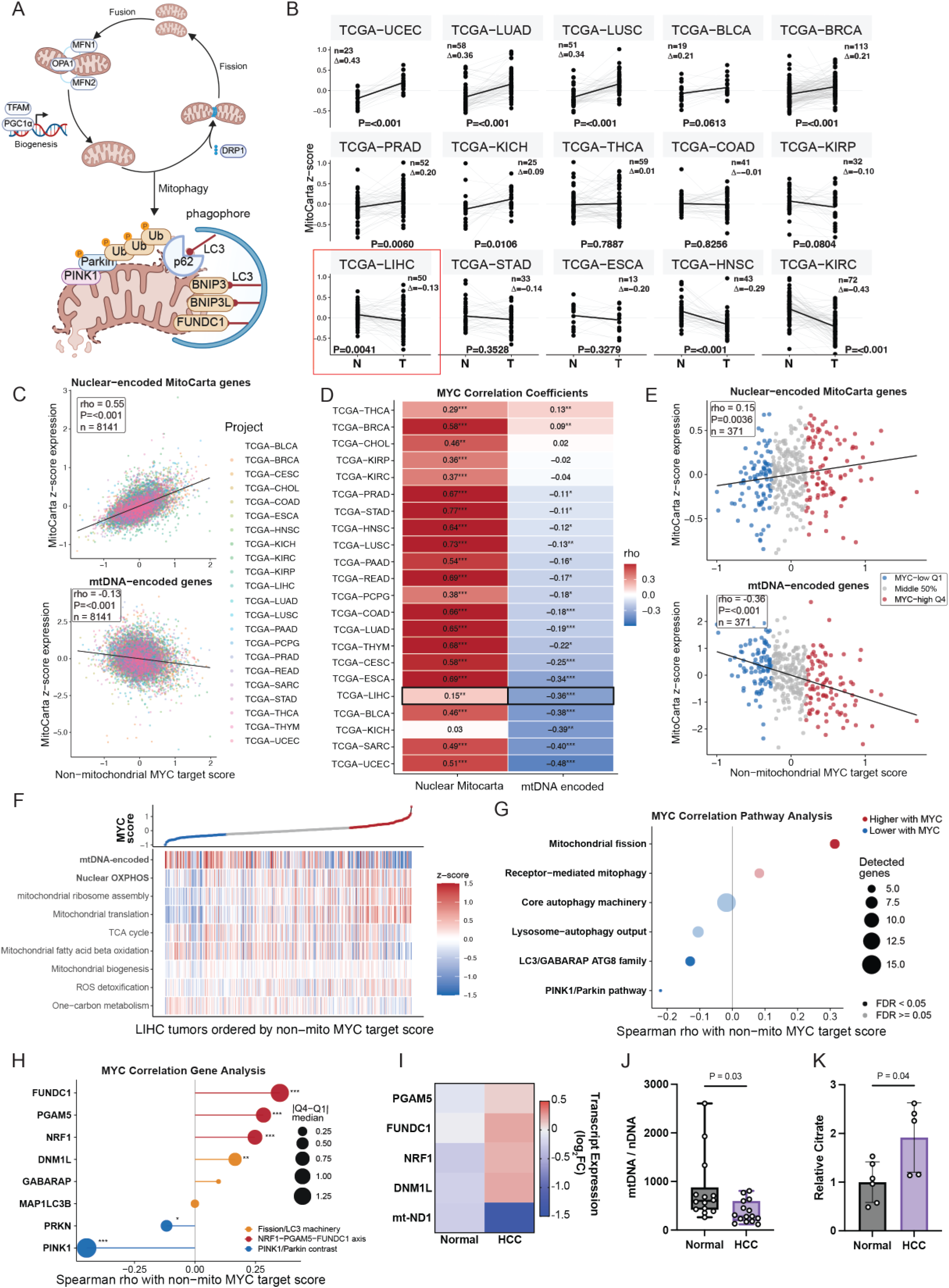
Pan-Cancer MYC–Mitochondrial Transcriptional Associations Converge on Mitophagy Dysregulation in Liver Cancer. **A)** Schematic detailing mitochondrial quality control (biogenesis, fission, fusion, and mitophagy) and the critical proteins. **B)** Paired MitoCarta gene expression z-scores in primary tumors (T) and adjacent normal tissue (N) across TCGA cancers. Lines connect matched samples, with black lines indicating cohort means; n, median tumor-minus-normal difference (Δ), and paired Wilcoxon signed-rank P values are shown. **C)** Pooled analysis of non-mitochondrial MYC target scores (a signature excluding mitochondria-localized targets) versus nuclear-encoded (top) and mtDNA-encoded (bottom) mitochondrial expression across TCGA primary tumors. Scores were standardized within cancer types before pooling; points are colored by cancer type, black lines indicate linear fits, and annotations report pooled Spearman rho, unadjusted P values, and sample sizes. **D)** Within-cancer Spearman correlations between non-mitochondrial MYC target scores and nuclear-encoded or mtDNA-encoded gene expression, across TCGA cancers. Tile colors and values indicate Spearman rho; asterisks denote Benjamini-Hochberg-adjusted significance (FDR < 0.05, 0.01, and 0.001 for *, **, and ***, respectively). **E)** Spearman correlations between non-mitochondrial MYC target scores and nuclear-encoded or mtDNA-encoded mitochondrial expression in TCGA-LIHC (n = 371). Black lines indicate linear fits; colors distinguish MYC-low (Q1), intermediate (Q2–Q3), and MYC-high (Q4) tumors. **F)** Heatmap of mitochondrial transcriptional program scores across TCGA-LIHC tumors ordered by increasing non-mitochondrial MYC target score. Colors indicate mean standardized expression, with MYC-low (Q1) and MYC-high (Q4) tumors highlighted in the upper track. **G)** Spearman correlations between non-mitochondrial MYC target scores and six curated mitophagy/autophagy programs. Dot size indicates the number of detected genes, color indicates correlation direction, and opacity distinguishes FDR-significant associations (Benjamini-Hochberg-adjusted P < 0.05). **H)** Spearman correlations between non-mitochondrial MYC target scores and individual mitochondrial quality-control genes. Dot size indicates the absolute difference in median standardized expression between MYC-high (Q4) and MYC-low (Q1) tumors; asterisks indicate Benjamini-Hochberg-adjusted P values: * < 0.05, ** < 0.01, and *** < 0.001. **I)** Heatmap of log_2_ RNA expression (normalized to actin) of mitochondrial quality control (*PGAM5, FUNDC1, NRF1, DNM1L* (DRP1)) and mtDNA-encoded gene (mt-ND1) in nonmatched, resected human tissue (n = 14 normal, n = 15 HCC). Color indicates increased (red) or decreased (blue) mean expression. **J)** Mitochondrial content (as defined by the ratio of mtDNA/nDNA) in nonmatched, human tissue (n = 14 normal, n = 15 HCC). Statistical significance was assessed by Welch’s two-tailed t test. **K)** Tumor citrate abundance relative to normal tissue as measured by GC-MS in a subset of nonmatched tissues collected with time-to-frozen <30 min (n = 6 normal, n = 5 HCC). Statistical significance was assessed by Welch’s two-tailed t test. TCGA study codes: BLCA-Bladder urothelial carcinoma; BRCA-Breast invasive carcinoma; CESC-Cervical squamous cell carcinoma and endocervical adenocarcinoma; CHOL-Cholangiocarcinoma; COAD-Colon adenocarcinoma; ESCA-Esophageal carcinoma; HNSC-Head and neck squamous cell carcinoma; KICH-Kidney chromophobe; KIRC-Kidney renal clear cell carcinoma; KIRP-Kidney renal papillary cell carcinoma; LIHC-Liver hepatocellular carcinoma; LUAD-Lung adenocarcinoma; LUSC-Lung squamous cell carcinoma; PAAD-Pancreatic adenocarcinoma; PCPG-Pheochromocytoma and Paraganglioma; PRAD-Prostate adenocarcinoma; READ-Rectum adenocarcinoma; SARC-Sarcoma; STAD-Stomach adenocarcinoma; THCA-Thyroid carcinoma; THYM-Thymoma; UCEC-Uterine corpus endometrial carcinoma

Mitophagy is regulated by multiple distinct mechanisms: the mitochondrial damage PINK1/Parkin pathway and the nutrient-sensitive BNIP3/NIX (BNIP3L) pathway.^11,12^ HCC often arises in the context of chronic liver disease and cirrhosis, associated with impaired hepatic circulation resulting in decreased oxygen (O_2_) availability.^13^ O_2_ tension regulates both OXPHOS activity and nutrient sensing, and several mitophagy pathways respond directly to low O_2_ availability through the transcription factor hypoxia-inducible factor 1 alpha (HIF-1α), frequently stabilized in HCC and associated with tumor progression.^14,15^ Specifically, Parkin inhibits HIF-1α stabilization, whereas BNIP3 and NIX are direct HIF-1α transcriptional targets, confirming the reciprocal relationship between mitophagy and O_2_ levels.^16,17^ An additional hypoxia-induced mitophagy receptor, FUN14 domain-containing 1 (FUNDC1), was shown to interact directly with microtubule-associated protein 1A/1B-light chain 3 (LC3) to stimulate mitophagy.^18^ This FUNDC1-LC3 interaction is regulated by phosphorylation events catalyzed by SRC and ULK1 kinases, as well as the phosphoglycerate mutase family member 5 (PGAM5) phosphatase.^19,20^ PGAM5 dephosphorylates FUNDC1 at Ser17, activating LC3 binding and mitophagy initiation. Together, these findings establish hypoxia as a key regulator of mitophagy pathway selection in the liver tumor microenvironment. Whether oncogene-specific signals selectively engage receptor-mediated versus damage-sensing mitophagy pathways in HCC remains unclear.

Altered mitochondrial quality control is implicated in HCC and other cancers. DRP1 has emerged as a promising target across cancers, especially in HCC, where it is often overexpressed.^10,21^ Some reports suggest this induction is amplified under hypoxia and that DRP1 inhibition delays tumor progression; however, others suggest that genetic DRP1 loss promotes spontaneous tumor formation and accelerated proliferation.^10,22,23^ Parkin-deficient mice are also susceptible to spontaneous liver tumor formation.^24^ Of note, BNIP3 is epigenetically silenced in human liver cancer patients, suggesting a potential tumor suppressive role for mitophagy.^25^ Furthermore, forced expression of mitochondrial fusion factors MFN2 or OPA1 in HCC results in apoptosis and is synergistic with BCL-2 inhibitors^26–28^. These conflicting results likely reflect the genetic heterogeneity of HCC, underscoring the need to examine mitochondrial quality control in the context of defined oncogenic drivers.

Here we show that MYC and β-catenin drive mechanistically distinct mitochondrial phenotypes in HCC. Contrary to MYC’s canonical role in promoting mitochondrial biogenesis, MYC-driven tumors paradoxically exhibit suppressed OXPHOS and reduced mitochondrial mass driven by elevated fission and receptor-mediated mitophagy through a *MYC-NRF1-FUNDC1/PGAM5/DNM1L*(DRP1) transcriptional axis, which reinforces glycolysis and anabolic metabolism. Genetic or pharmacologic inhibition of this pathway reduced mitophagy, restored oxidative phosphorylation, and extended survival in tumor-bearing mice. Critically, restoration of mitochondrial function alone, independently of mitophagy, was sufficient to reduce tumor burden. These results demonstrate that mitochondrial suppression is an actively maintained tumor-promoting dependency and identify mitochondrial quality control as a therapeutic vulnerability in MYC-driven HCC.

## Results

### MYC Activity in Human HCC Correlates with Mitophagy Signatures

Mitochondrial remodeling is a common feature of cancer; however, whether distinct oncogenic drivers alter mitochondrial transcriptional programs in similar ways remains unclear. To define how mitochondrial gene expression changes during tumorigenesis, we first compared mitochondrial gene expression in matched tumor and adjacent normal tissues across TCGA cohorts using MitoCarta 3.0 (a curated, experimentally validated dataset of 1,136 human genes encoding mitochondrially localized proteins) to quantify mitochondrion-related transcriptional output. Paired analysis revealed that mitochondrial gene expression is tumor-type specific, as it is elevated in some cancers (e.g., UCEC, LUAD, BRCA) but suppressed in others, including liver hepatocellular carcinoma (LIHC) (**Figure 1B, Figure S1A**).

As *MYC* activation is a recurrent feature of LIHC and other aggressive tumor types and tightly linked to cellular metabolism, we next asked whether increased MYC transcriptional activity correlates with changes in mitochondrial gene expression. We excluded all MitoCarta-defined mitochondrial genes from the Hallmark MYC Targets V1 and V2 gene sets before scoring (**Figure S1B**). We confirmed that this refined MYC activity score (hereafter ‘non-mito MYC target score’) captures genuine MYC transcriptional output (**Figure S1C, D**). Paired tumor-normal analysis demonstrated that total MYC target gene expression increased in all tumor types relative to adjacent normal tissue (**Figure S1E, F**), consistent with broad *MYC* activation during tumorigenesis.

We then examined whether MYC activity was associated with mitochondrial gene transcriptional output across cancers. Mitochondrial function depends on gene products encoded by both the nuclear and mitochondrial genomes, yet these transcript classes are often considered together. We therefore stratified mitochondrial genes by their genomic origin (nuclear-encoded versus mtDNA-encoded) to determine whether mitochondrial programs were concordantly or discordantly regulated by MYC activity. MYC-mitochondrial transcriptional discordance was observed broadly across tumor types (**Figure 1C, D**), with nuclear-encoded transcripts moderately and positively correlating with MYC activity (rho = 0.55) while mtDNA-encoded transcripts were weakly and inversely correlated (rho = −0.13). This discordance was particularly accentuated in liver cancer, where the positive association between MYC and nuclear-encoded transcripts was attenuated (rho = 0.19) while the inverse correlation with mtDNA-encoded transcripts was markedly stronger (rho = −0.32). The discordance was also amplified in *MYC*-high tumors (**Figure 1E, F**). Disparate regulation contrasts with MYC-associated mitochondrial biogenesis and prompted us to hypothesize that selective suppression of mtDNA-encoded transcripts reflects altered mitochondrial turnover in MYC-driven HCC.

One mechanism that could selectively deplete mtDNA-encoded transcripts while preserving nuclear-encoded ones is mitochondrial quality control. We therefore asked whether MYC activity correlated with mitochondrial dynamics and mitophagy transcriptional signatures (**Figure 1G**). Strong associations were detected between *MYC* activity and *FUNDC1*-, *PGAM5*-, *NRF1*-, and *DNM1L*(DRP1)-associated pathways, whereas canonical *PINK1*/*PARKIN* pathway genes showed a negative association with MYC activity (**Figure 1H**). PINK1/PARKIN-mediated mitophagy is canonically activated by overt mitochondrial damage and membrane depolarization, whereas FUNDC1/PGAM5/NRF1-mediated mitophagy is nutrient-and hypoxia-sensitive, suggesting that MYC preferentially engages stress-responsive rather than damage-responsive mitophagy programs.

We confirmed these results in an independent cohort (n=14 normal, n=15 HCC), demonstrating that *MYC* expression is positively associated with *FUNDC1, PGAM5, NRF1,* and *DNM1L* (**Figure S1G**) and each was upregulated in HCC compared to normal liver tissue (**Figure 1I**). Furthermore, tissue from this cohort displayed decreased expression of the mtDNA-encoded *ND1* gene (**Figure 1I**) and reduced mitochondrial content (**Figure 1J**). Consistent with impaired mitochondrial respiratory function, HCC tissues exhibited elevated citrate levels (**Figure 1K**), indicative of TCA cycle stalling and reduced OXPHOS flux. Notably, patients with increased *MYC*, mitophagy, and *NRF1*, *FUNDC1*, *PGAM5*, and *DNM1L* signatures exhibit significantly worse survival outcomes (**Figure S2**).

Together, these analyses identify a conserved *MYC*-associated mitochondrial state in HCC characterized by discordance between nuclear-and mtDNA-encoded mitochondrial gene transcription, and consistent with elevated receptor-mediated mitophagy and mitochondrial remodeling. To test this hypothesis *in vivo,* we employed murine HCC models associated with oncogenic *MYC* signaling.

### MYC and β-catenin Drive Distinct Mitochondrial Phenotypes in HCC

These TCGA results were surprising given the well-known associations between MYC and mitochondrial biogenesis.^6^ Although HCC tumors are genetically heterogeneous, *MYC* and *CTNNB1* are among the most common oncogenic drivers (observed in 18% and 27% of human HCC, respectively) (**Figure 2A**). We employed the sleeping beauty transposon transposase hydrodynamic tail vein injection (SBTT-HDTVI) model of HCC^5^ (**Figure 2B**) to examine how these oncogenes influence HCC progression and metabolism. Control mice received hydrodynamic injection of a transposon plasmid lacking an oncogene insert (hereafter ‘Empty’) to provide non-tumor liver tissue for comparison. β-catenin-driven tumors were also selected as a comparator because β-catenin represents a prevalent HCC oncogenic driver, produces a well-differentiated tumor morphology that is distinct from MYC-driven tumors, and shares the PTEN loss background, allowing oncogene-specific metabolic effects to be isolated (**Figure 2C**). All models result in rapid lethality and allow investigation into oncogene-specific effects while avoiding alterations from chronic liver damage (**Figure 2D**).

**Figure 2.**
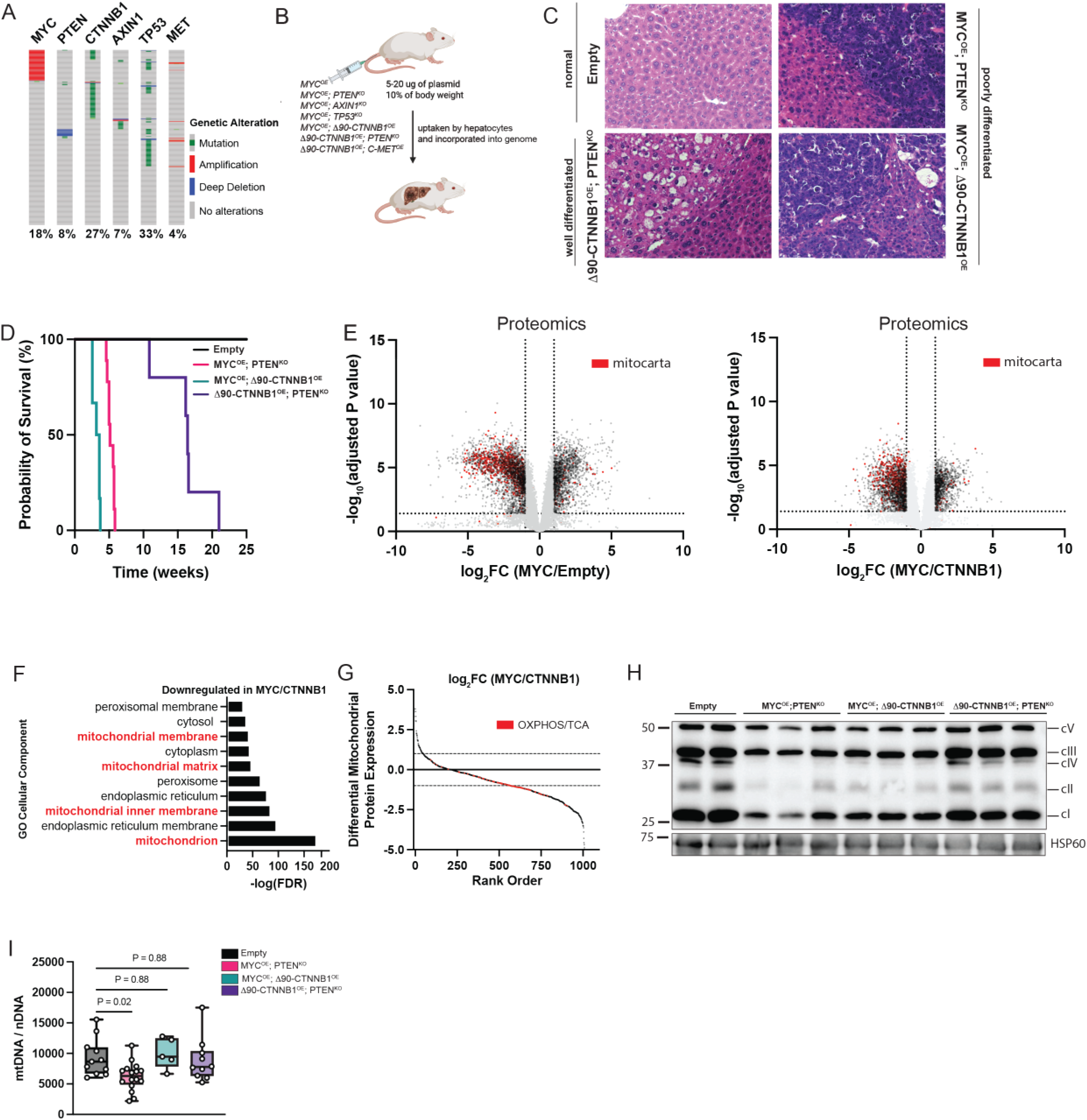
MYC-Driven HCC Exhibits Selective Downregulation of Mitochondrial Proteins and Oxidative Phosphorylation. **A)** OncoPrint of frequency and type of genetic alterations (mutation [green], amplification [red], or deletion [blue]) in selected oncogenes (*MYC, PTEN, CTNNB1, AXIN1, TP53, MET*) used in this study from TCGA-LIHC (n=366). **B)** Schematic of the hydrodynamic tail vein injection model used to deliver Sleeping Beauty transposon and transposase constructs encoding the indicated oncogenes into mouse hepatocytes to induce HCC. Oncogene combinations are listed in the figure. **C)** Representative H&E histology of HCC tumors showing poorly differentiated MYC tumors (*MYC^OE^; PTEN^KO^, MYC^OE^; CTNNB1^OE^*) and well-differentiated β-catenin tumors (*CTNNB1^OE^; PTEN^KO^*) (n=5 per group). Scale bar = 100 um. OE = overexpression; KO = knockout. **D)** Kaplan-Meier survival curves for HCC-bearing mice of the indicated oncogene combinations (panel C, n=5-8 per group). Endpoint was defined when mice showed signs of morbidity and/or a >15% weight gain. **E)** Volcano plot of protein abundances in *MYC^OE^; PTEN^KO^* tumors compared to empty control liver or *MYC^OE^; PTEN^KO^* tumors compared to *CTNNB1^OE^; PTEN^KO^* tumors, based on label-free proteomics quantification. Log_2_(Fold change) is plotted against the –log_10_(adjusted p-value) for each protein. Significantly changing proteins (log_2_(Fold change)>1 or < –1; Adjusted p-val <0.05) are colored black, and significantly altered mitochondrial proteins (Mitocarta) are colored red (n=4-6 per group). **F)** Bar chart of DAVID GO Cellular Component enrichment analysis of significantly downregulated proteins (log_2_FC < –1, adjusted P < 0.05) in *MYC^OE^; PTEN^KO^* compared to *CTNNB1^OE^; PTEN^KO^* tumors. The top 10 enriched terms are shown; bar length indicates – log10(FDR). All shown terms meet P < 0.05. **G)** Rank order of average log_2_ fold change of all mitochondrial proteins in *MYC^OE^; PTEN^KO^*tumors versus *CTNNB1^OE^; PTEN^KO^*. Selected OXPHOS and TCA Cycle proteins are colored red. **H)** Representative western blot of OXPHOS proteins (cV – ATP5A; cIII – UQCRC2; cIV – mtCO1; cII – SDHB; cI – NDUFB8) of the indicated genotype (Empty, *MYC^OE^; PTEN^KO^, MYC^OE^; CTNNB1^OE^,* and *CTNNB1^OE^; PTEN^KO^)* (n=2-3 biological replicates per group). HSP60 was used as a loading control. **I)** Mitochondrial content (as defined by the mtDNA / nDNA ratio) across the indicated genotypes (n=5-7 per group). Statistical significance was assessed using two-tailed t test with Dunnett correction for multiple comparisons.

We performed label-free proteomics (**Table S1**) on HCC tumors and control livers (**Figure 2E**) and reanalyzed RNA-sequencing data from Molina-Sanchez et al., 2020 (GSE148379, **Table S2, S3A-B**). To confirm that these models represent transcriptomically, proteomically, and metabolomically distinct tumor subtypes, we performed principal component analysis (PCA) across platforms (**Figure S3C-E**), which demonstrated clear separation between MYC and β-catenin tumors. Both proteomics and RNA-sequencing demonstrated significant reductions in mitochondrially localized proteins and transcripts, respectively (**Figure 2E, S3A-B**). DAVID (Database for Annotation, Visualization, and Integrated Discovery) functional annotation on significantly downregulated genes in *MYC^OE^*; *PTEN^KO^* vs *CTNNB1^OE^*; *PTEN^KO^* tumors (hereafter referred to as MYC tumors and β-catenin tumors, respectively) indicated that mitochondrial processes were significantly disrupted (**Figure 2F, G**). Loss of OXPHOS proteins was observed across all HCC tumors but was most pronounced in MYC tumors (**Figure 2H, Table S1**). We confirmed that OXPHOS loss occurred in MYC tumors regardless of secondary oncogenic driver and was independent of mouse strain (FVB and C57/B6) (**Figure S3F-H**). Consistent with these molecular phenotypes, mitochondrial content was significantly decreased in MYC tumors compared to control liver tissue and β-catenin driven tumors (**Figure 2I**). *MYC^OE^; CTNNB1^OE^* tumors produced severe lethality and heterogeneous molecular phenotypes, with individual tumors resembling either MYC-or β-catenin-driven tumors depending on which oncogenic program predominated. Multi-omic profiling revealed platform-specific clustering: transcriptomic profiles were distinct from both MYC and β-catenin-driven models, proteomic profiles clustered adjacent to β-catenin tumors, and metabolomic profiles fell intermediate between the two (**Figure S3C–E**), consistent with context-dependent oncogene dominance rather than a fixed co-operative transcriptional program. Together, the distinct and reproducible phenotypes of MYC vs β-catenin models recapitulate those observed in human HCC and provide genetically defined systems for mechanistic investigation.

Having established that MYC and β-catenin tumors represent distinct molecular subtypes, we next sought to define the metabolic pathways that most distinguish them. Joint pathway analysis combining log_2_ fold changes in proteins and metabolites identified pathways that span cytosolic and mitochondrial compartments, including pyruvate metabolism, the TCA cycle, and urea cycle/arginine metabolism, as well as glycolysis (**Figure 3A-B**). Suppressed urea cycle (UC) metabolism is a defining feature of HCC, and UC activity was lost across all tumors consistent with mRNA, protein, and metabolite alterations (**Figure S4A-D**). Decreased UC activity (**Figure 3C**) was confirmed directly using an ornithine transcarbamylase (OTC) activity assay; OTC is the key enzyme that condenses carbamyl phosphate and ornithine to produce citrulline. Glutamate dehydrogenase (GDH) and glutamine synthetase (GLUL) activity were only mildly perturbed (**Figure S4E, F**), suggesting that alternative ammonia clearance pathways were insufficient to compensate for UC loss, thereby contributing to increased circulating ammonia (**Figure 3D**). This was compounded by decreased abundance of the ammonia exporter, *SLC42A2* (RHBG) (**Figure S4G**), and increased intratumoral ammonia in MYC tumors (**Figure 3E**). Isotopic tracing with [^15^NH_4_Cl] confirmed that tumor-bearing mice converted less ammonia into urea with no compensatory glutamate or glutamine labeling (**Figure 3F, G**), demonstrating a functional defect in nitrogen metabolism.

**Figure 3.**
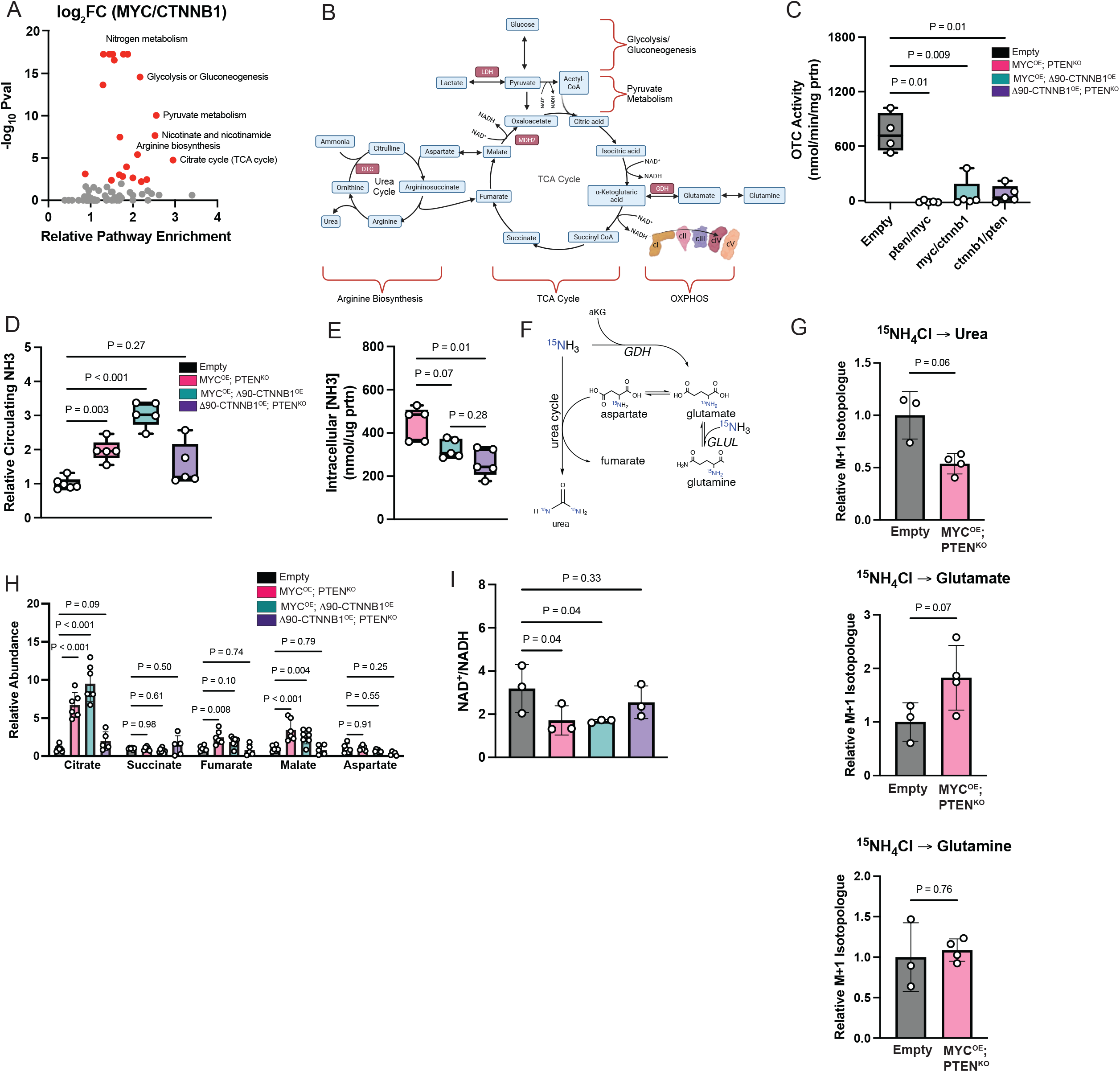
HCC Oncogene-Dependent Mitochondrial Metabolic Reprogramming is Characterized by Ammonia Accumulation and TCA Cycle Dysregulation. **A)** Bubble plot of MetaboAnalyst pathway enrichment analysis comparing *MYC^OE^; PTEN^KO^* to *CTNNB1^OE^; PTEN^KO^* tumors, using combined metabolomic and proteomic log_2_ fold changes as input (n=5–7 per group). The x-axis indicates pathway impact score and the y-axis indicates – log10(P value). Significantly (Holm adjusted p-value <0.05) altered pathways are highlighted red; top enriched pathways are labeled. ***B)*** Integrated schematic of top metabolic pathways (arginine biosynthesis, TCA Cycle, OXPHOS, pyruvate metabolism, glycolysis/gluconeogenesis) identified in pathway analysis (A). **C)** *Ex-vivo* ornithine transcarbamylase (OTC) activity (nmol/min/mg protein) in snap-frozen liver or tumor tissue of the indicated genotype (n=4-5 per group). Genotypes were as follows: Empty, *MYC^OE^; PTEN^KO^, MYC^OE^; CTNNB1^OE^*; *CTNNB1^OE^; PTEN^KO^*. Statistical significance was assessed by one-way Welch’s ANOVA with Dunnett’s T3 post-hoc test for multiple comparisons against the empty vector control. **D)** Enzymatic assay of plasma ammonia in empty (no tumor) or tumor-bearing mice (n=5-6 per group). Empty mice were normalized to one. Genotypes were as follows: Empty, *MYC^OE^; PTEN^KO^, MYC^OE^; CTNNB1^OE^*; *CTNNB1^OE^; PTEN^KO^*. Statistical significance was assessed by one-way Welch’s ANOVA with Dunnett’s T3 post-hoc test for multiple comparisons against the empty vector control. **E)** Intracellular ammonia concentration (nmol/protein) in tumor tissue as measured through a GDH enzymatic assay (n=4-5 per group). Genotypes were as follows: *MYC^OE^; PTEN^KO^, MYC^OE^; CTNNB1^OE^*; *CTNNB1^OE^; PTEN^KO^*. Statistical significance was assessed by one-way Welch’s ANOVA with Dunnett’s T3 post-hoc test for multiple comparisons across all groups. **F)** Schematic of ^15^NH_4_Cl intracellular labeling fates (urea cycle, glutamate dehydrogenase (GDH), or glutamine synthetase (GLUL)). **G)** Relative M+1 isotopologue abundance of urea, glutamate, and glutamine in empty or *MYC^OE^; PTEN^KO^* tissue 5 minutes post ^15^NH_4_Cl injection (5mmol/kg) as measured by GC-MS (n=3-4 per group). Statistical significance was assessed using two-tailed Welch’s t test. **H)** TIC-normalized GC-MS TCA cycle metabolite (citrate, succinate, fumarate, malate, and aspartate) levels in tumors relative to empty tissue (n=5-7 per group). Genotypes were as follows: Empty, *MYC^OE^; PTEN^KO^, MYC^OE^; CTNNB1^OE^*; *CTNNB1^OE^; PTEN^KO^*. Statistical significance was assessed by one-way Welch’s ANOVA with Dunnett’s T3 post-hoc test for multiple comparisons against the empty vector control. **I)** Enzymatic quantification of redox ratio (NAD^+^/NADH) in tumors (n=3 per group). Genotypes were as follows: Empty, *MYC^OE^; PTEN^KO^, MYC^OE^; CTNNB1^OE^*; *CTNNB1^OE^; PTEN^KO^*. Statistical significance was assessed by one-way Welch’s ANOVA with Dunnett’s T3 post-hoc test for multiple comparisons against the empty vector control.

Although increased metabolic recycling of ammonia into glutamate was not observed (**Figure 3G**), uncleared ammonia can act as a mitochondrial uncoupler.^29^ *Ex vivo* ammonia treatment decreased the respiratory control ratio (**Figure S4H-J**), and MYC-derived cell lines showed greater respiratory sensitivity to ammonia than β-catenin cells (**Figure S4K**). MYC tumors displayed significant TCA cycle stalling with accumulation of fumarate, malate, and citrate (**Figure 3H**) and a decreased NAD^+^/NADH ratio (**Figure 3I**), confirming that suppressed OXPHOS is a defining feature of MYC-driven HCC, further exacerbated by hyperammonemia.

To establish whether these phenotypes reflect MYC activity directly, as opposed to ammonia accumulation or differentiation status, we performed three independent tests. A low-protein diet reduced ammonia but did not affect OXPHOS protein levels or survival (**Figure S5A-C**) in tumor-bearing mice. Wildtype mice fed high ammonium acetate chow showed no reduction in OXPHOS proteins or TCA cycle disruption (**Figure S5D-I**). Furthermore, *MYC^OE^; AXIN1^KO^* mice, which produce well-differentiated tumors with normal circulating ammonia levels, fully recapitulated the MYC mitochondrial phenotype, including OXPHOS loss and TCA stalling (**Figure S6**). Together, these results confirm that MYC is the primary driver of mitochondrial dysfunction in HCC, with hyperammonemia as a compounding but not causative factor.

### MYC Drives Oxidative Stress and Mitochondrial Fragmentation

Consistent with impaired OXPHOS flux, MYC tumors displayed elevated intracellular ROS (**Figure 4A**) and total glutathione (**Figure 4B),** suggestive of compensatory upregulation of glutathione synthesis. We measured phosphorylated histone H2AX (γH2AX pS139) levels to assess DNA damage as a functional readout of oxidative stress, and only MYC tumors displayed significant DNA damage (**Figure 4C**).

**Figure 4.**
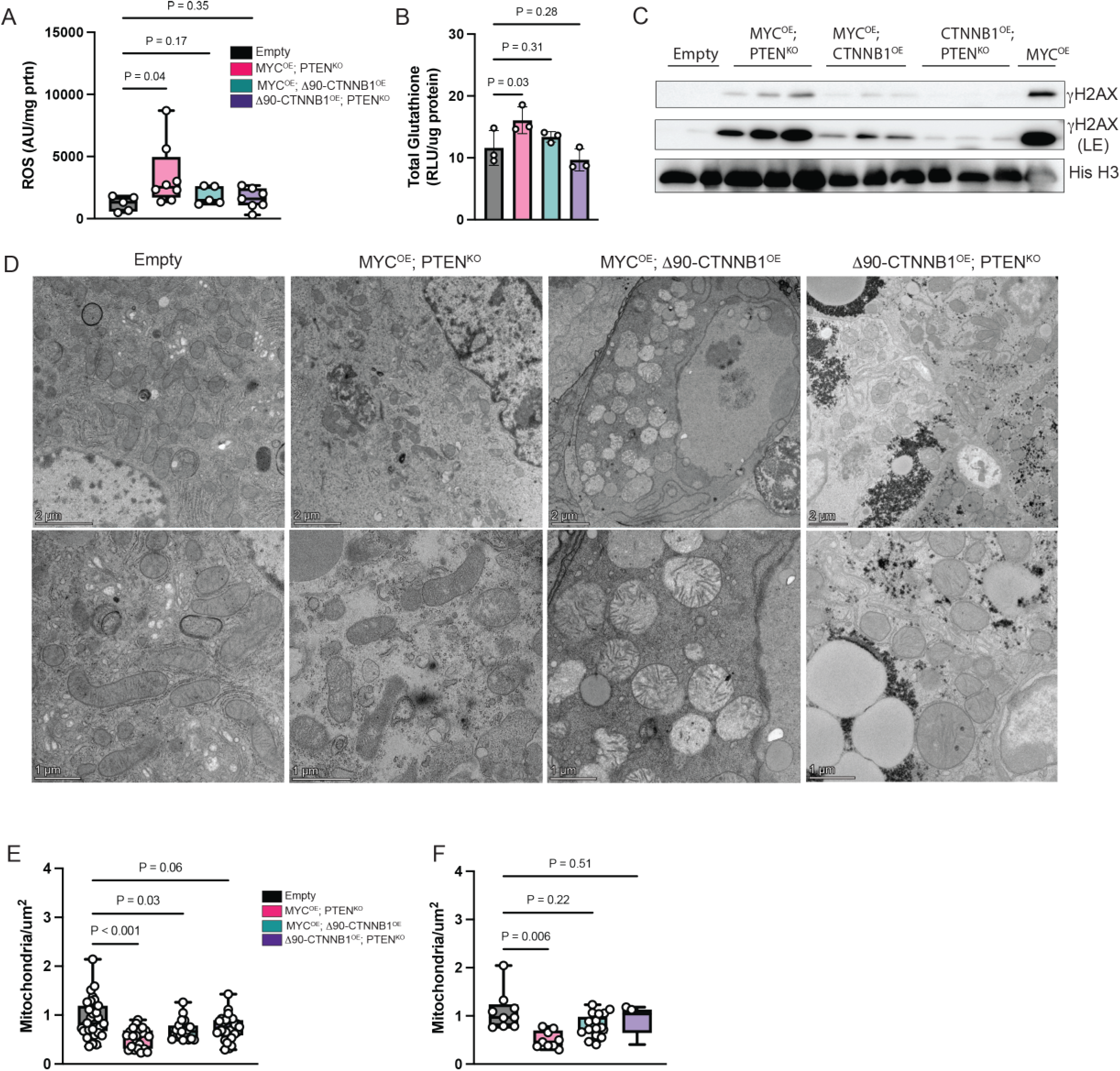
Oxidative Stress and Mitochondrial Remodeling Are Hallmarks of MYC-Driven HCC **A)** Luciferase-based ROS-Glo H_2_O_2_ levels normalized to protein (RLU/ mg protein) from snap-frozen tissue (n=5-8 per group). Genotypes were as follows: Empty, *MYC^OE^; PTEN^KO^, MYC^OE^; CTNNB1^OE^*; *CTNNB1^OE^; PTEN^KO^*. Statistical significance was assessed by one-way Welch’s ANOVA with Dunnett’s T3 post-hoc test for multiple comparisons against the empty vector control. **B)** Enzymatic quantification of total glutathione (RLU/ mg protein) in tissue normalized to protein (n=3 per group). Genotypes were as follows: Empty, *MYC^OE^; PTEN^KO^, MYC^OE^; CTNNB1^OE^*; *CTNNB1^OE^; PTEN^KO^*. Statistical significance was assessed by one-way Welch’s ANOVA with Dunnett’s T3 post-hoc test for multiple comparisons against the empty vector control. **C)** Representative DNA damage in tissue as determined by γH2AX (pS139) western blotting (n=2-3 per group). Genotypes were as follows: Empty, *MYC^OE^; PTEN^KO^, MYC^OE^; CTNNB1^OE^*; *CTNNB1^OE^; PTEN^KO^*. Histone H3 was used as a loading control. This panel is representative and not quantified. **D)** Representative TEM images of mitochondrial morphology in tumor tissue of the indicated genotypes. Genotypes were as follows: Empty, *MYC^OE^; PTEN^KO^, MYC^OE^; CTNNB1^OE^*; *CTNNB1^OE^; PTEN^KO^*(representative of n=3 biological replicates). Top panels show overview morphology (scale bar = 2 um) and bottom panels show higher magnification of mitochondria structure and cristae (scale bar = 1um). **E)** Quantification of mitochondrial length (um) in TEM. Mitochondrial length was measured from (n=10 mitochondria/image per biological replicate (n=3)) using ImageJ. Genotypes were as follows: Empty, *MYC^OE^; PTEN^KO^, MYC^OE^; CTNNB1^OE^*; *CTNNB1^OE^; PTEN^KO^*. Statistical significance was assessed by one-way Welch’s ANOVA with Dunnett’s T3 post-hoc test for multiple comparisons against the empty vector control. **F)** TEM quantification of number of mitochondria per area (mitochondria/um^2^). Mitochondria per area was measured by counting the number of mitochondria per image (n=3 image per biological replicate (n=3)) using ImageJ. Images were acquired at 4300X and area was calculated from the scale bar. Genotypes were as follows: Empty, *MYC^OE^; PTEN^KO^, MYC^OE^; CTNNB1^OE^*; *CTNNB1^OE^; PTEN^KO^*. Statistical significance was assessed by one-way Welch’s ANOVA with Dunnett’s T3 post-hoc test for multiple comparisons against the empty vector control.

Given the observed loss of OXPHOS proteins, TCA cycle metabolite accumulation, and increased ROS, we next performed transmission electron microscopy (TEM) to determine whether these functional deficits were accompanied by structural remodeling of the mitochondria. Whereas MYC tumors displayed significantly smaller mitochondria and fewer mitochondria per unit area, β-catenin tumors displayed significant mitochondrial swelling, with irregular cristae, shortened length, but relatively preserved mitochondrial number per area (**Figure 4D-F**). The small, circular mitochondria in MYC tumors were consistent with mitochondrial fragmentation, whereas the swelling observed in β-catenin tumors may reflect a mechanistically distinct form of mitochondrial dysfunction that does not converge on fission-driven mitophagy.

Collectively, these data demonstrate that MYC tumors exhibit a signature of oxidative stress, DNA damage, and mitochondrial disruption distinct from the mitochondrial swelling observed in β-catenin tumors. Decreased mitochondria and mitochondrial fragmentation in MYC tumors suggest increased mitophagic flux, which we next tested through genetic perturbation of mitophagy.

### Receptor-Mediated Mitophagy Suppresses OXPHOS and Drives Tumor Progression

Consistent with our TCGA analysis (**Figure 1**), compared to β-catenin tumors, murine MYC tumors upregulated multiple components of the receptor-mediated mitophagy machinery (**Figure 5A),** with specific increases in NRF1, PGAM5, FUNDC1, and p-FUNDC1 (Ser17), a direct ULK1 target.^30^ Prior work has shown that FUNDC1 and PGAM5 cooperate to activate mitophagy downstream of *NRF1* and *MYC.*^30^ To assess the functional importance of this receptor-mediated mitophagy pharmacologically, we treated mice with the ULK1 inhibitor SBI-0206965 (50mg/kg, every 3 days), which suppresses both general autophagy and mitophagy. Pharmacologic inhibition extended mouse survival by approximately 25% (**Figure 5B**) and increased mitochondrial respiration (**Figure 5C**), confirming that mitophagy inhibition and associated restoration of mitochondrial function promote mouse survival. Having established pharmacological proof-of-concept, we next asked which specific receptor pathway was most critical.

**Figure 5.**
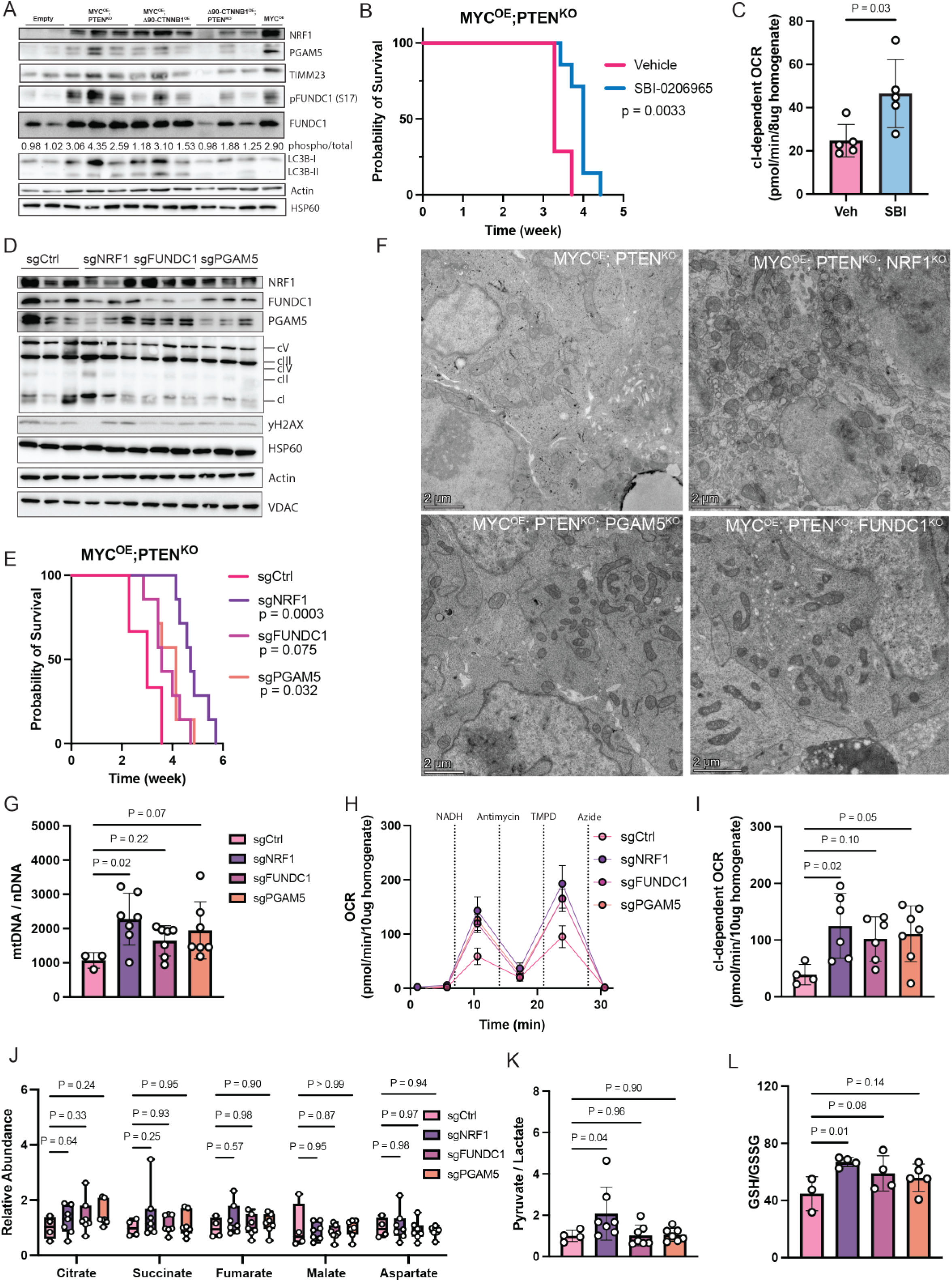
Mitophagy Inhibition and Restoration of OXPHOS Delay Tumor Progression. **A)** Representative western blot analysis of mitophagy proteins (NRF1, PGAM5, FUNDC1 (total and pSer17), LC3B-I, and LC3B-II) identified in TCGA-LIHC transcriptomic data (Figure 1). Actin and HSP60 were used as loading controls (n=2-3 per group). Genotypes were as follows: Empty, *MYC^OE^; PTEN^KO^, MYC^OE^; CTNNB1^OE^*; *CTNNB1^OE^; PTEN^KO^*. This panel is representative and not quantified. **B)** Kaplan-Meier survival curve with log-rank (Mantel-Cox) test of *MYC^OE^; PTEN^KO^*mice treated every 3 days with ULK1 inhibitor SBI-0206965 or vehicle (IP injection 50mg/kg) starting 2 weeks post oncogene delivery until endpoint (n=7 per group). P-values are displayed in the figure. Endpoint was defined when mice showed signs of morbidity and/or a >15% weight gain. **C)** Seahorse analysis of complex I-dependent (NADH) mitochondrial respiration (pmol/min/ ug protein) from frozen *MYC^OE^; PTEN^KO^* tumor tissue treated with vehicle or SBI-0206965 (n = 5 per group). Complex I-dependent was determined by the NADH – Antimycin A OCR. Statistical significance was assessed using two-tailed Welch’s t test. **D)** Representative western blot analysis showing NRF1, FUNDC1, and PGAM5 CRISPR/Cas9 knockout efficiency. OXPHOS (cV – ATP5A; cIII – UQCRC2; cIV – mtCO1; cII – SDHB; cI – NDUFB8), and γH2AX (pS139) were also determined. HSP60 (mitochondrial matrix), Actin (whole cell), and VDAC (mitochondrial outer membrane) were used as loading controls (n=3 per group). This blot is representative and not quantified. Genotypes were as follows: *MYC^OE^; PTEN^KO^* (sgCtrl*), MYC^OE^; PTEN^KO^; NRF1^KO^* (sgNRF1), *MYC^OE^; PTEN^KO^; FUNDC1^KO^* (sgFUNDC1*), MYC^OE^; PTEN^KO^; PGAM5^KO^* (sgPGAM5). **E)** Kaplan-Meier survival curve with pairwise log-rank (Mantel-Cox) test of *MYC^OE^; PTEN^KO^* (sgCtrl)vs. *MYC^OE^; PTEN^KO^; NRF1^KO^* (sgNRF1) vs. *MYC^OE^; PTEN^KO^; FUNDC1^KO^* (sgFUNDC1*), vs. MYC^OE^; PTEN^KO^; PGAM5^KO^* (sgPGAM5) mice (n=7-8 per group). P-values are displayed in the figure. Color scheme is maintained throughout subsequent panels sgCtrl (black), sgNRF1 (purple), sgFUNDC1 (pink), sgPGAM5 (orange). Endpoint was defined when mice showed signs of morbidity and/or a >15% weight gain. **F)** Representative TEM images of mitochondrial morphology and abundance in tumor tissue of the indicated genotypes (representative of n=3 biological replicates). Genotypes were as follows: *MYC^OE^; PTEN^KO^* (sgCtrl*), MYC^OE^; PTEN^KO^; NRF1^KO^* (sgNRF1), *MYC^OE^; PTEN^KO^; FUNDC1^KO^* (sgFUNDC1*), MYC^OE^; PTEN^KO^; PGAM5^KO^* (sgPGAM5). Panels show overview morphology (scale bar = 2 um). **G)** Mitochondrial content (as defined by the ratio of mtDNA/nDNA) in mitophagy knockout genotypes (n = 3-7 per group). Genotypes were as follows: *MYC^OE^; PTEN^KO^* (sgCtrl*), MYC^OE^; PTEN^KO^; NRF1^KO^* (sgNRF1), *MYC^OE^; PTEN^KO^; FUNDC1^KO^* (sgFUNDC1*), MYC^OE^; PTEN^KO^; PGAM5^KO^* (sgPGAM5). Statistical significance was assessed by one-way Welch’s ANOVA with Dunnett’s T3 post-hoc test for multiple comparisons against the sgCtrl. **H)** Seahorse analysis from frozen tissue homogenates given complex I (NADH) and complex IV (TMPD/Ascorbate) substrates (n=4-7 per group). Genotypes were as follows: *MYC^OE^; PTEN^KO^* (sgCtrl*), MYC^OE^; PTEN^KO^; NRF1^KO^* (sgNRF1), *MYC^OE^; PTEN^KO^; FUNDC1^KO^* (sgFUNDC1*), MYC^OE^; PTEN^KO^; PGAM5^KO^* (sgPGAM5). Representative OCR (pmol/min/ug protein) trace is shown. **I)** Complex I-dependent (NADH) respiration (pmol/min/ug protein) in frozen tissue homogenates (n = 4-7 per group). Complex I-dependent was determined by the NADH – Antimycin A OCR. Genotypes were as follows: *MYC^OE^; PTEN^KO^* (sgCtrl*), MYC^OE^; PTEN^KO^; NRF1^KO^* (sgNRF1), *MYC^OE^; PTEN^KO^; FUNDC1^KO^* (sgFUNDC1*), MYC^OE^; PTEN^KO^; PGAM5^KO^* (sgPGAM5). Statistical significance was assessed by one-way Welch’s ANOVA with Dunnett’s T3 post-hoc test for multiple comparisons against the sgCtrl. **J)** TIC-normalized GC-MS TCA cycle (citrate, succinate, fumarate, malate, and aspartate) metabolites in knockout tumors (n = 4-7 per group). Genotypes were as follows: *MYC^OE^; PTEN^KO^* (sgCtrl*), MYC^OE^; PTEN^KO^; NRF1^KO^* (sgNRF1), *MYC^OE^; PTEN^KO^; FUNDC1^KO^* (sgFUNDC1*), MYC^OE^; PTEN^KO^; PGAM5^KO^* (sgPGAM5). Statistical significance was assessed by one-way Welch’s ANOVA with Dunnett’s T3 post-hoc test for multiple comparisons against the sgCtrl. **K)** Pyruvate / lactate ratio (proxy of cytosolic NAD^+^/NADH ratio) from GC-MS analysis (n = 4-7 per group). Genotypes were as follows: *MYC^OE^; PTEN^KO^* (sgCtrl*), MYC^OE^; PTEN^KO^; NRF1^KO^* (sgNRF1), *MYC^OE^; PTEN^KO^; FUNDC1^KO^* (sgFUNDC1*), MYC^OE^; PTEN^KO^; PGAM5^KO^* (sgPGAM5). Statistical significance was assessed by one-way Welch’s ANOVA with Dunnett’s T3 post-hoc test for multiple comparisons against the sgCtrl. **L)** Luciferase-based GSH/ GSSG-Glo ratio as a marker of oxidative stress in tumors (n=3-5 per group). Genotypes were as follows: *MYC^OE^; PTEN^KO^* (sgCtrl*), MYC^OE^; PTEN^KO^; NRF1^KO^* (sgNRF1), *MYC^OE^; PTEN^KO^; FUNDC1^KO^* (sgFUNDC1*), MYC^OE^; PTEN^KO^; PGAM5^KO^* (sgPGAM5). Statistical significance was assessed by one-way Welch’s ANOVA with Dunnett’s T3 post-hoc test for multiple comparisons against the sgCtrl.

To determine how individual mitophagy receptor pathways control survival in the context of MYC-driven HCC, we generated *NRF1*, *PGAM5*, and *FUNDC1* knockout tumors in the *MYC^OE^; PTEN^KO^* background using SBTT-HDTVI, with knockout efficiency confirmed by western blot (**Figure 5D**). *FUNDC1* knockout resulted in a non-significant (p = 0.075) increase in survival, while *PGAM5* and *NRF1* knockouts significantly increased survival by 38% and 57%, respectively (**Figure 5E**). This graded survival benefit is consistent with NRF1 occupying a master regulatory position upstream of both PGAM5 and FUNDC1. TEM demonstrated increased mitochondrial number and electron density in knockout tumors (**Figure 5F**), reflecting accumulation of metabolically active mitochondria. Mechanistically, loss of receptor machinery prevented mitochondrial recognition and clearance, leading to the accumulation of fragmented mitochondria.

Consistent with TEM observations, tumors demonstrated an increase in mitochondrial content (**Figure 5G**) and enhanced complex I-dependent respiration (**Figure 5H, I**). TCA cycle metabolites remained unchanged (**Figure 5J**), suggesting that OXPHOS restoration is reflected in redox balance. Consistent with this, OXPHOS suppression forces cells to rely on lactate dehydrogenase for NAD^+^ regeneration, thereby preferentially shifting pyruvate toward lactate and suppressing the pyruvate-to-lactate ratio. Restoration of complex I respiration would be expected to relieve this glycolytic pressure, and we observed increased pyruvate to lactate ratios (representing the cytosolic NAD^+^/NADH) in knockout tumors (**Figure 5K**). This shift confirms that the Warburg-like shift in MYC-driven HCC is a functional consequence of OXPHOS suppression rather than primary transcriptional reprogramming. The restoration of respiratory efficiency in receptor knockout tumors decreased cytosolic cytochrome c (**Figure S7A**), indicative of decreased apoptosis and increased mitochondrial fidelity, and resolved oxidative stress (**Figure 5L**). These results suggest that mitophagy programs drive tumor progression in MYC-driven HCC by suppressing mitochondrial function and disrupting redox homeostasis.

### DRP1 Promotes HCC Progression via Mitochondrial Fission and ROS Generation

Mitophagy knockouts resulted in accumulation of fragmented mitochondria; therefore, we sought to understand the role of mitochondrial fission in the observed phenotypes. Pharmacologic inhibition of DRP1 using the heptapeptide P110 was well-tolerated (**Figure S7B**), decreased tumor burden after 5 weeks (**Figure 6A**), and resulted in a non-significant (p = 0.085) increase in mouse survival (**Figure S7C**). We then knocked out *DNM1L* (DRP1) in *MYC^OE^; PTEN^KO^* tumors using SBTT-HDTVI. *DRP1* knockout mice displayed increased survival (p = 0.0002) compared to wildtype mice (**Figure 6B)**, although DRP1 showed only partial knockout by RNA (**Figure 6C**) and protein (**Figure 6D**). Consistent with prior reports^23^, mice with complete *DRP1* loss developed hepatic adenomas, whereas tumors retaining partial DRP1 expression showed greater malignant transformation (**Figure S7D)**. These results likely reflect transient Cas9 expression in the SBTT system and selection for tumor cells retaining partial DRP1 activity and collectively underscore the complexity of DRP1’s function.

**Figure 6.**
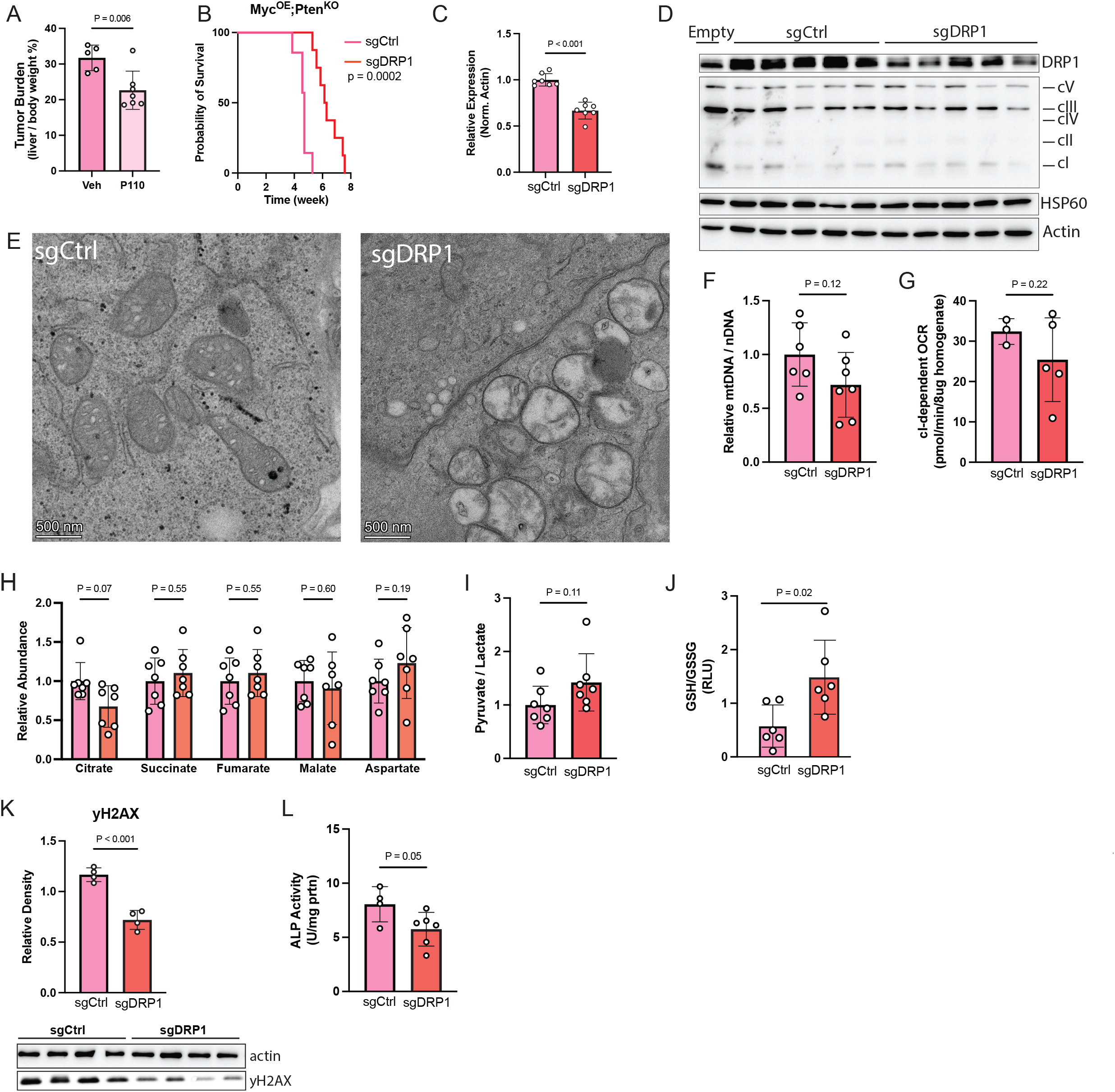
DRP1 Promotes Tumor Growth Through ROS Generation and Mitophagy Activation. **A)** Tumor burden (liver weight / body weight) in *MYC^OE^; PTEN^KO^* mice treated with DRP1 inhibitor (a DRP1-specific inhibitory peptide, TAT-P110) or vehicle (TAT) via IP injections (5mg/kg/day) starting two weeks post oncogene delivery until 5 weeks post oncogene injection (n=5-7 per group). Statistical significance was assessed using two-tailed Welch’s t test. **B)** Kaplan-Meier survival curve with log-rank (Mantel-Cox) test of *MYC^OE^; PTEN^KO^; DNM1L^WT^* (sgCtrl) vs. *MYC^OE^; PTEN^KO^; DNM1L^KO^* (sgDRP1) mice (n=7-8 per group). P-values are displayed in the figure. Color scheme is maintained throughout subsequent panels: sgCtrl (pink) and sgDRP1 (red). Endpoint was defined when mice showed signs of morbidity and/or a >15% weight gain. **C)** Relative mRNA expression normalized to actin of DRP1 in tumors of the indicated genotypes as measured by qRT-PCR (n=7 per group). Genotypes were as follows: *MYC^OE^; PTEN^KO^; DNM1L^WT^* (sgCtrl) vs. *MYC^OE^; PTEN^KO^; DNM1L^KO^* (sgDRP1). Statistical significance was assessed using two-tailed Welch’s t test. **D)** Representative western blot analysis of DRP1 and OXPHOS (cV – ATP5A; cIII – UQCRC2; cIV – mtCO1; cII – SDHB; cI – NDUFB8) proteins (n=5 per group). HSP60 and Actin were used as loading controls. This blot is representative and not quantified. Genotypes were as follows: *MYC^OE^; PTEN^KO^; DNM1L^WT^* (sgCtrl) vs. *MYC^OE^; PTEN^KO^; DNM1L^KO^* (sgDRP1). **E)** Representative TEM images (n=3 biological replicates) of tumor mitochondria swelling and cristae disruption. Scale bar = 500 nm. Genotypes were as follows: *MYC^OE^; PTEN^KO^; DNM1L^WT^* (sgCtrl) vs. *MYC^OE^; PTEN^KO^; DNM1L^KO^* (sgDRP1). **F)** Mitochondrial content (as defined by the ratio of mtDNA/nDNA) of tumors (n=6-7 per group). Genotypes were as follows: *MYC^OE^; PTEN^KO^; DNM1L^WT^* (sgCtrl) vs. *MYC^OE^; PTEN^KO^; DNM1L^KO^* (sgDRP1). Statistical significance was assessed using two-tailed Welch’s t test. **G)** Complex I-dependent (NADH) respiration (pmol/min/ug protein) in frozen tumor tissue homogenates (n = 3-5 per group). Complex I-dependent was determined by the NADH – Antimycin A OCR. Genotypes were as follows: *MYC^OE^; PTEN^KO^; DNM1L^WT^* (sgCtrl) vs. *MYC^OE^; PTEN^KO^; DNM1L^KO^* (sgDRP1). Statistical significance was assessed using two-tailed Welch’s t test. **H)** TIC normalized, GC-MS TCA cycle metabolite (citrate, succinate, fumarate, malate, aspartate) abundance in tumors (n=7 per group). Genotypes were as follows: *MYC^OE^; PTEN^KO^; DNM1L^WT^* (sgCtrl) vs. *MYC^OE^; PTEN^KO^; DNM1L^KO^* (sgDRP1). Statistical significance was assessed using two-tailed Welch’s t test. **I)** Pyruvate / lactate ratio (proxy of cytosolic NAD^+^/NADH ratio) from GC-MS analysis (n = 7 per group). Genotypes were as follows: *MYC^OE^; PTEN^KO^; DNM1L^WT^(sgCtrl)* vs. *MYC^OE^; PTEN^KO^; DNM1L^KO^*(sgDRP1). Statistical significance was assessed using two-tailed Welch’s t test. **J)** Luciferase-based GSH/ GSSG-Glo ratio as a marker of oxidative stress in tumors (n=6 per group). Genotypes were as follows: *MYC^OE^; PTEN^KO^; DNM1L^WT^* (sgCtrl) vs. *MYC^OE^; PTEN^KO^; DNM1L^KO^* (sgDRP1). Statistical significance was assessed using two-tailed Welch’s t test. **K)** DNA damage quantified by yH2AX (pS139) western blot (n=4 per group) as a marker of DNA damage (densitometry normalized to Actin using ImageJ). Actin was used as a loading control. Genotypes were as follows: *MYC^OE^; PTEN^KO^; DNM1L^WT^* (sgCtrl) vs. *MYC^OE^; PTEN^KO^; DNM1L^KO^* (sgDRP1). Statistical significance was assessed using two-tailed Welch’s t test. **L)** Colorimetric alkaline phosphatase (proxy for lysosomal degradative capacity) in tumors normalized to protein (IU/ mg protein) (n= 4-6 per group). Genotypes were as follows: *MYC^OE^; PTEN^KO^; DNM1L^WT^* (sgCtrl) vs. *MYC^OE^; PTEN^KO^; DNM1L^KO^* (sgDRP1). Statistical significance was assessed using two-tailed Welch’s t test.

TEM confirmed the structural remodeling, with DRP1 knockout tumors displaying increased mitochondrial size and swelling (**Figure 6E**), resembling the morphology of β-catenin driven tumors, which do not activate DRP1. While DRP1 loss improved survival, it did not rescue OXPHOS proteins (**Figure 6D**), mitochondrial content (**Figure 6F**), respiration (**Figure 6G**), TCA cycle metabolite levels (**Figure 6H**), or the NAD^+^/NADH ratio (**Figure 6I, S7E**) compared to *MYC^OE^; PTEN^KO^* tumors. Instead, DRP1 loss significantly reduced oxidative stress (**Figure 6J, K, S7F**), likely through restoration of mitochondrial GSH import.^31^ Despite structural remodeling and reduced fission substrates, DRP1 knockout tumors displayed a modest decrease in alkaline phosphatase (ALP) activity (**Figure 6L**), indicating that lysosomal-autophagic flux (and mitophagy) was not fully abolished. This is consistent with prior work demonstrating that while DRP1-mediated fission promotes the generation of fragmented mitochondria as the canonical substrate for mitophagy, receptor-mediated mitophagy is not strictly dependent on DRP1-driven fragmentation.^32–34^ Together, our mitophagy and DRP1 knockout results confirm that residual receptor-mediated mitophagy is sufficient to maintain OXPHOS suppression in the absence of fission and implicate downstream mitophagy receptors as the primary executors of mitochondrial clearance in MYC-driven HCC. These findings demonstrate why inhibiting fission is insufficient to fully restore respiratory capacity and highlight mitophagy receptors as a primary therapeutic target.

### OXPHOS Restoration is Sufficient to Delay Tumor Progression

Having established that receptor-mediated mitophagy maintains OXPHOS suppression, we asked whether directly restoring OXPHOS, independently of mitophagy, is sufficient to suppress tumor growth. To test this, we generated tumors containing yeast NADH dehydrogenase (NDI1) or a catalytically inactive mutant (NDI1-E272A) as a control. NDI1 bypasses mitochondrial complex I to directly oxidize NADH and donate electrons to coenzyme Q and is rotenone resistant (**Figure 7A**). NDI1 activity was confirmed by measurement of rotenone-resistant respiration (**Figure 7B)**. At 5 weeks post-injection, when tumors are established but mice are not at endpoint, NDI1-expressing mice showed reduced tumor burden compared to NDI1-E272A controls (**Figure 7C**), demonstrating that restoration of mitochondrial electron transport is sufficient to suppress tumor growth.

**Figure 7.**
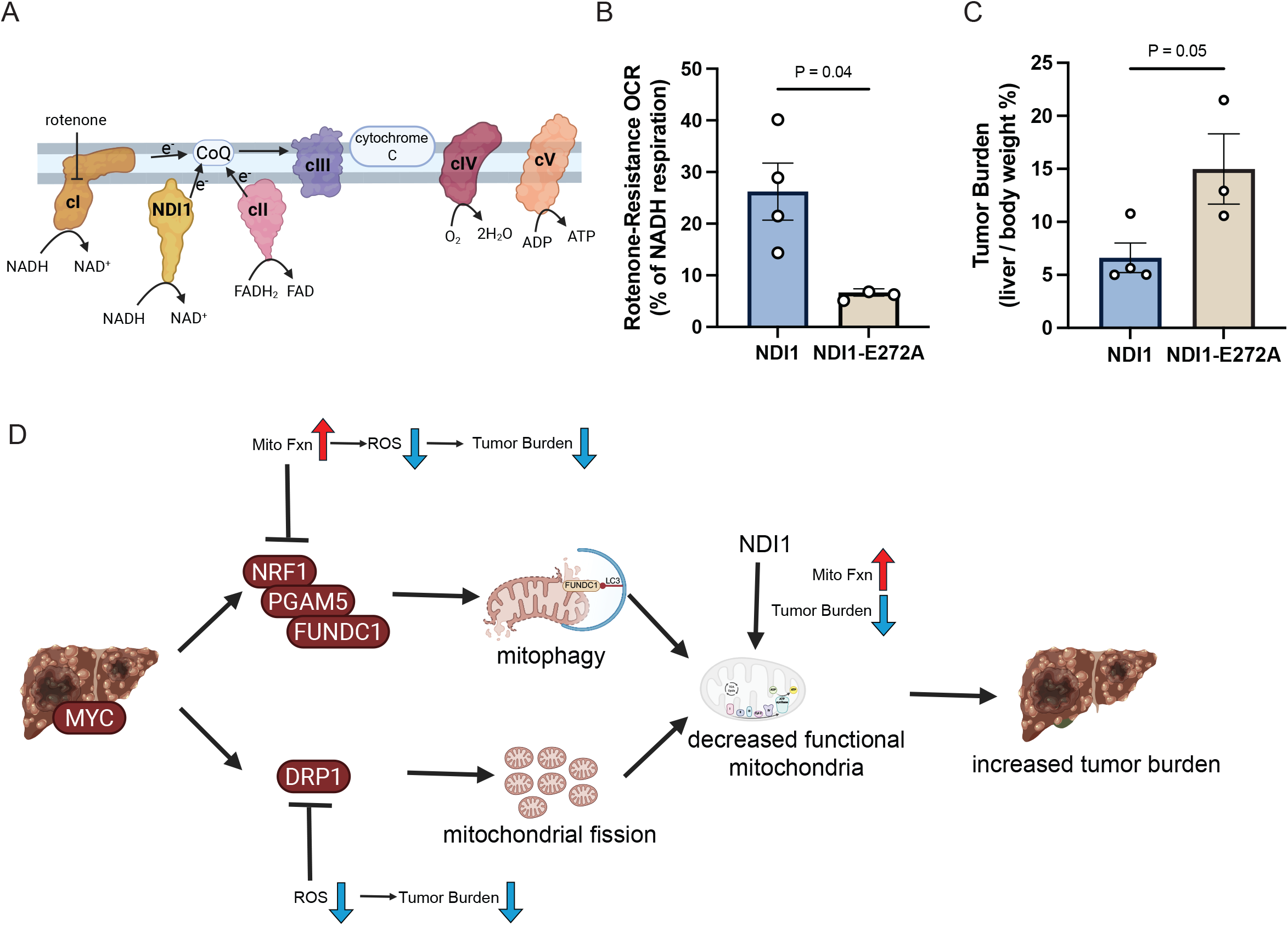
Bypassing Complex I Restores Mitochondrial Respiration and Suppresses Tumor Growth, Identifying the MYC-NRF1-Mitophagy Axis as a Therapeutic Vulnerability in HCC. **A)** Schematic illustrating the mechanism by which yeast NDI1, a rotenone-insensitive NADH dehydrogenase, bypasses mammalian complex I to donate electrons directly to coenzyme Q (CoQ) and restore OXPHOS activity. NDI1-E272A, a catalytically inactive point mutant, serves as a negative control. **B)** Rotenone-resistant complex I respiration (OCR in the presence of rotenone; pmol/min/ug protein, a measurement of NDI1 activity) in *MYC^OE^; PTEN^KO^; NDI1^OE^* (NDI1) and *MYC^OE^; PTEN^KO^; NDI1^E272A^* (NDI1-E272A, catalytically inactive control) frozen tumors by Seahorse (n=3-4 per group). Statistical significance was assessed using two-tailed Welch’s t test. **C)** Tumor burden (liver weight / body weight) in *MYC^OE^; PTEN^KO^; NDI1^OE^* (NDI1) and *MYC^OE^; PTEN^KO^; NDI1^E272A^* (NDI1-E272A, catalytically inactive control) 5 weeks post oncogene delivery (n = 3-4 per group). Statistical significance was assessed using two-tailed Welch’s t test. **D)** Proposed model of MYC-NRF1-mitophagy axis and suppression of OXPHOS in HCC. MYC overexpression drives the NRF1-PGAM5-FUNDC1 axis, which promotes mitophagy and decreases mitochondrial function and increases tumor burden. Inhibition of NRF1-PGAM5-FUNDC1 axis increases mitochondrial function, decreasing ROS and tumor burden. DRP1 inhibition also decreases ROS, thereby reducing tumor burden through a distinct mechanism. Rescue of dysfunctional OXPHOS through NDI1 expression restores mitochondrial function and reduces tumor burden.

Together, these data establish that MYC-driven HCC suppresses OXPHOS through a DRP1-initiated, receptor-mediated mitophagy program in which NRF1 acts as the central transcriptional coordinator (**Figure 7D**). Suppression of OXPHOS by this pathway forces cellular reliance on glycolysis for NAD^+^ regeneration, creating a state of metabolic inflexibility that sustains tumor growth. Blocking this pathway at the level of the fission machinery partially impairs tumor progression by reducing ROS and restoring mitochondrial GSH import.

Meanwhile, receptor-mediated mitophagy inhibition achieves a more complete reversal by restoring mitochondrial respiratory capacity, resolving redox imbalance, and relieving glycolytic dependence, thereby reducing tumor burden. Direct OXPHOS restoration through NDI1 is sufficient to recapitulate these antitumor effects. Our results establish enhanced mitochondrial electron transport as a key tumor-suppressive mechanism and provide preclinical rationale for targeting the MYC-mitophagy-OXPHOS axis in patients with MYC-driven hepatocellular carcinoma.

## Discussion

We demonstrate that MYC suppresses mitochondrial function through elevated *NRF1*-driven mitophagy, uncoupled from its canonical role in biogenesis (**Figure 1, S1**). In MYC-dependent liver cancer, mitochondrial abundance and function are decreased, resulting in increased ROS levels, elevated mitochondrial fission, and upregulated receptor-mediated mitophagy to promote tumor cell proliferation. Inhibiting mitophagy through genetic or pharmacologic mechanisms extended the survival of tumor-bearing mice by reducing ROS levels and restoring mitochondrial function (**Figure 7D**). Bypassing complex I with NDI1 reduced tumor burden independently of mitophagy, confirming that OXPHOS suppression is a maintained tumor dependency. These results, combined with clinical data, suggest that activation of mitochondrial quality control pathways decreases patient survival and represents an important metabolic vulnerability in HCC (**Figure S2**).

As noted above, we observed a paradoxical inverse relationship between *MYC* activity and OXPHOS gene expression, which we propose reflects the interplay between *MYC* and hypoxia signaling in the tumor microenvironment. Although *MYC* is known to promote mitochondrial biogenesis through its transcriptional target genes *TFAM* and *PGC1α*, and shift cellular metabolism towards aerobic glycolysis (the Warburg effect), mitochondrial intermediates are still required to produce biosynthetic substrates. HCC tumors are significantly hypoxic with a median pO_2_ as low as 0.8%.^35^ The interplay between HIF-α and MYC is complex and has not been fully characterized in the context of the high-level MYC amplification typical of MYC-driven HCC.^36^ Hypoxia can suppress respiration and mitochondrial biogenesis, partly through HIF-α.^36,37^ Consistent with our hypothesis, we observed that tumors with increased HIF-1α protein abundance negatively correlate with ETC protein abundance (**Table S2**). *HIF* signaling can further suppress MYC transcriptional activity through multiple mechanisms, including MAX competition and *MXI1* induction, compounding the suppression of mitochondrial biogenesis.^38^

We propose the following model: under hypoxia, decreased ETC activity and elevated ROS reduce mitochondrial membrane potential, triggering DRP1 recruitment and mitophagy. Simultaneously, HIF-α activation inhibits mitochondrial biogenesis, compounding the net loss of mitochondrial mass over the course of tumor progression. This model aligns with the proteomics and transcriptomics data (**Table S1, S2**), neither of which revealed increased TFAM or PGC1-α abundance, consistent with decreased biogenesis. Over the course of tumor progression, this interplay results in a significant decrease in mitochondrial mass, explaining reduced mitochondrial content, decreased OXPHOS capacity, and suppressed mtDNA-encoded transcripts.

A key mechanistic finding is that receptor-mediated mitophagy through FUNDC1, PGAM5, and NRF1 sustains mitochondrial clearance and OXPHOS suppression independently of DRP1-driven fission. Although DRP1 fragmentation generates canonical mitophagy substrates, autophagosomes can still engage receptor machinery and clear mitochondria in its absence. As a result, DRP1 loss decreased oxidative stress, but did not restore OXPHOS. DRP1 deficient mitochondria remained structurally swollen with irregular cristae yet still showed reduced oxidative stress, suggesting that the act of fission is a primary ROS driver.

Receptor-mediated mitophagy knockout tumors reduced ROS through a mechanistically distinct route, accumulating functional mitochondria with restored ETC capacity that reduces electron leak through more efficient coupling of electron transfer in the respiratory chain. Strikingly, DRP1 knockout and β-catenin tumors displayed convergent mitochondrial morphology, both showing large, swollen mitochondria, reflecting their shared absence of active DRP1-mediated fission. This convergence revealed that the relatively preserved mitochondrial function of β-catenin tumors is the result of an inactive receptor-mediated mitophagy, identifying this pathway as a targetable dependency specific to MYC-driven HCC.

Mitophagy-driven OXPHOS suppression has consequences extending beyond respiratory capacity. When OXPHOS proteins are cleared by mitophagy, cells become dependent on cytosolic lactate dehydrogenase for NAD^+^ regeneration and sustain glycolytic flux. This glycolytic dependence represents a state of metabolic inflexibility: tumor cells can no longer utilize oxidative metabolism to meet biosynthetic or energetic demands, and instead become committed to a highly-glycolytic, high-cytosolic NADH state. Restoration of complex I-dependent respiration in receptor knockout tumors relieves this pressure by recovering oxidative NAD^+^ regeneration, confirming that the Warburg-like metabolic shift in MYC-driven HCC is a functional consequence of mitophagy-driven OXPHOS suppression. Shifting the redox balance toward mitochondrial NAD^+^ oxidation (via NDI1 expression) would be expected to reduce carbon flux into reductive biosynthetic pathways, restricting the anabolic substrates available to support tumor growth.

Mitochondrial function is critical for cellular proliferation,^39^ and as a result, OXPHOS has been identified as a therapeutic target. The mitochondrial complex I inhibitor, IACS-010759, showed promising pre-clinical inhibition of OXPHOS and delayed tumor growth in multiple cancers, including HCC.^40^ However, clinical trials were discontinued due to peripheral neuropathy, a common phenotype in mitochondrial disorders.^41^ Conversely, mitochondrial dysfunction is considered a core feature in cancers, suggesting that activation of OXPHOS is a mechanism to promote apoptosis in cancer.^42^ Increasing OXPHOS by inhibiting pyruvate dehydrogenase kinase (PDK) promotes apoptosis by increasing ROS levels, a mechanism critical to chemoresistance.^43^ OXPHOS suppression reinforces glycolysis and reductive biosynthesis, whereas OXPHOS restoration shifts NADH oxidation to the mitochondria. Our findings suggest that targeting mitochondrial quality control, rather than directly inhibiting complex I, may offer a more tumor-selective therapeutic window in HCC that promotes OXPHOS. Collectively, we suggest that the metabolic state imposed by MYC-driven mitophagy (low OXPHOS, high cytosolic NADH, elevated glycolysis) represents an actively maintained dependency that can be reversed to suppress tumor growth.

Oncogene-dependent mitochondrial phenotypes in HCC are clinically significant. A recent prospective study demonstrated subtype-specific differences in response to transarterial chemoembolization (TACE), the most widely used locoregional therapy for unresectable HCC. Weinfurtner et al.^44^ biopsied 53 HCC tumors prior to TACE and found that tumors harboring activating Wnt/β-catenin pathway mutations achieved significantly higher rates of complete response (54.5% vs. 22.6%, p=0.02) and longer time to tumor progression (not reached vs. 8.3 months, p=0.02). Mechanistic studies revealed that β-catenin mutant HCC cells depend on OXPHOS for ATP production and are unable to compensate via glycolytic ATP generation under ischemic conditions. Our findings illuminate the molecular underpinning of this clinical observation: β-catenin-driven HCC preserves mitochondrial mass and OXPHOS capacity, which is exploited by TACE-induced ischemia. In contrast, MYC-driven HCC, in which the *NRF1-PGAM5-FUNDC1* mitophagy axis actively suppresses OXPHOS and enforces a Warburg-like glycolytic state, would be predicted to tolerate ischemia and exhibit reduced TACE sensitivity. This suggests the *NRF1-PGAM5-FUNDC1* mitophagy axis functions as a tumor-intrinsic metabolic regulator and as a determinant of locoregional therapy response, with implications for biomarker-guided treatment selection. Together, these results suggest that restoring OXPHOS capacity (through mitophagy inhibition) could sensitize MYC-driven HCC tumors to TACE.

Our study focused on primary HCC; however, the role of OXPHOS and TCA cycle activity in HCC metastasis remains unaddressed. Metastatic tumors in other cancer types often show increased OXPHOS relative to primary tumors, and it will be important to determine whether the mitochondrial dysfunction observed here restricts or enables metastatic spread.^45,46^ Given that HCC most commonly metastasizes via the portal vein,^47^ defining this metabolic adaptation could have implications for managing post-transplant relapse. Additionally, the impact of altered mitochondrial function on the tumor immune microenvironment, particularly given that mitochondrial ROS regulates innate immune sensing, warrants future investigation. Finally, although our pharmacologic studies demonstrate proof-of-concept for targeting receptor-mediated mitophagy, the development of either NRF1-selective inhibitors or receptor-specific mitophagy antagonists optimized for hepatic delivery represents an important translational priority for MYC-amplified HCC patients, who currently have no approved targeted therapy.

## Methods

### MICE

All experiments and procedures were reviewed and approved by the Institutional Animal Care and Use Committee (IACUC, protocol 804389) at the University of Pennsylvania. Mice were housed in a controlled environment (12 h light/12 h dark cycle, humidity 30-70%, temperature 20-22 °C) and had free access to water and rodent diet. All animal experiments were performed in accordance with the Guide for Care and Use of Laboratory Animals of the NIH.

Sleeping Beauty Transposon Transposase hydrodynamic tail vein injections were performed on 6 – 8 week old male FVB/NJ (Jax #001800) or C57/B6 (Jax #000664) mice. Briefly, mice were injected with transposase pCMV(CAT)T7-SB100, transposons c-myc-PT3EF1a, pT3-N90-beta-catenin, pT3-EF1a-c-Met, and Cas9 plasmids: px330-Pten-2, px330-Axin1, or px330 p53 in the indicated combinations. For dual-knockout experiments an sgPTEN sequence and sgDRP1, sgPGAM5, sgFUNDC1, or sgNRF1 sequence was cloned into px333. For overexpression experiments c-MYC and an internal ribosome entry site (IRES) was expressed with *Saccharomyces cerevisiae* NADH-ubiquinone reductase (NDI1) (Vectorbuilder VB260501-1222qgy) or mt-Keima (Vectorbuilder VB260605-1634tww). NDI1-E272A was generated using Q5 site-directed mutagenesis kit (NEB #E0554S) with the following primers (F – TCACTTGGTCgcaGCTCTGCCCA and R – ATTTGGACTTCTTCGGCCAATG)

Plasmids were injected at a 5:1 ratio of transposon: transposase. Plasmids were diluted in sterile saline at a final volume of 10% of the mouse’s weight (20 – 25g) and injected over 5-8 seconds. Mice were allowed to recover for 30 minutes before being returned to their cage. Plasmids were acquired from Addgene. pCMV(CAT)T7-SB100 was a gift from Zsuzsanna Izsvak (Addgene plasmid # 34879; http://n2t.net/addgene:34879; RRID:Addgene_34879). c-myc-PT3EF1a was a gift from Xin Chen (Addgene plasmid # 92046; http://n2t.net/addgene:92046; RRID:Addgene_92046). pT3-N90-beta-catenin was a gift from Xin Chen (Addgene plasmid # 31785; http://n2t.net/addgene:31785; RRID:Addgene_31785). px330-Pten-2 was a gift from Amaia Lujambio (Addgene plasmid #162552; http://n2t.net/addgene:162552; RRID:Addgene_162552). pX330 p53 was a gift from Tyler Jacks (Addgene plasmid # 59910; http://n2t.net/addgene:59910; RRID:Addgene_59910). pX333 was a gift from Andrea Ventura (Addgene plasmid # 64073; http://n2t.net/addgene:64073; RRID:Addgene_64073). pT3-EF1a-c-Met was a gift from Xin Chen (Addgene plasmid # 31784; http://n2t.net/addgene:31784; RRID:Addgene_31784).

Mice used in the low protein diet experiment were injected and then divided into two groups. One group was fed a 20% protein control diet (Picolab Rodent Diet 5053), while the other was given a 6% protein diet (Teklad TD.90016).

TAT-P110 (MCE #HY-P2280) was dissolved in saline and injected IP daily at 5mg/kg starting 2 weeks after HDTVI. A control TAT peptide (MCE #HY-P0281) was used as the vehicle control. SBI-0206965 (MCE #HY-16966) was dissolved in 10% DMSO, 40% PEG400, 5% Tween80, and 45% saline and IP injected every 3 days at 50mg/kg. ^15^NH^4^Cl was IP injected into tumor bearing mice at 5mmol/kg and animal sacrificed 5 minutes post injection. All animals were sacrificed by cervical dislocation and tissue collected and frozen immediately for downstream processing.

### Western Blot

Tissue was crushed on liquid nitrogen. To roughly 10 mg of tissue, 10 volumes of lysis buffer [50 mM Tris pH 7.6, 2% SDS, 1X Halt protease and phosphatase inhibitor (Thermo Scientific #78441)] were added. Samples lysed on ice for 5 – 10 minutes, then centrifuged at max speed for 10 minutes at 4 °C. Protein content was measured by BCA assay (Thermo Scientific #23225). Primary antibodies used: Total-OXPHOS rodent (Abcam #ab110413), DRP1 (Cell Signaling Technology #5391), HSP60 (Cell Signaling Technology #1265), VDAC (Cell Signaling Technology #4661), Histone H3 (Cell Signaling Technology #4499), phospho-Histone H2A.X (S139) (Cell Signaling Technology #9718), OPA1 (Cell Signaling Technology #80471), Mitofusin-2 (Cell Signaling Technology #9482), B Actin (Cell Signaling Technology #4970), GLUD1 (Proteintech #14299), CPS1 (Proteintech #18703), Pyruvate Carboxylase (ABclonal #A6301), HSP90 (Cell Signaling Technology #4874), ASS1 (Cell Signaling Technology #70720), Argininosuccinate Lyase (Novus Biologicals #NBP1-32752), OTC (Abcam #ab203859), Arg1 (Cell Signaling Technology #93668), GLUL (Proteintech #11037), B Tubulin (Cell Signaling Technology #2146), NRF1 (Cell Signaling Technology #46743), PGAM5 (Cell Signaling Technology #24584), TIMM23 (Cell Signaling Technology #34822), FUNDC1 (Cell Signaling Technology #49240), phospho-FUNDC1 (S17) (Thermo Scientific #PA5-114576). All primary antibodies were used at 1:1000 diluted in 5% bovine serum albumin. Secondary antibodies used: Anti-rabbit IgG, HRP-linked (Cell Signaling Technology #7074) and Anti-mouse IgG, HRP-linked (Cell Signaling Technology #7076).

### Gene Expression Analyses

RNA was isolated from crushed tissue using the RNEasy Mini kit (Qiagen #74104). RNA was converted to cDNA (Fisher Scientific #4387406) for qPCR. The following primers were used: CPS1 (Thermo Scientific #Mm01256489), OTC (Thermo Scientific #Mm00493267), ASS1 (Thermo Scientific #Mm00711256), ASL (Thermo Scientific #Mm01197741), Arg1 (Thermo Scientific #Mm00475988), Arg2 (Thermo Scientific #Mm00477592), NAGS (Thermo Scientific #Mm00467530), SLC7A1 (Thermo Scientific #Mm01219063), GLUD1 (Thermo Scientific #Mm00492353), GLUL (Thermo Scientific #Mm00725701), DNM1L (Thermo Scientific #Mm01342903).

### TCGA transcriptomic and pan-cancer analyses

GDC-processed STAR gene counts for TCGA cohorts were obtained from the NCI Genomic Data Commons using TCGAbiolinks. Primary tumor and solid-tissue-normal samples were identified from TCGA sample-type barcodes. Matched tumor-normal samples were used for paired analyses, whereas all eligible primary tumors were included in tumor-only analyses. Cancer types were processed independently.

Low-expression genes were filtered before normalization and variance-stabilizing transformation with DESeq2. Within each cohort, expression was standardized for each gene across samples, and gene-set scores were calculated as the mean standardized expression of detected member genes.

Mitochondrial genes were defined using Human MitoCarta3.0 (https://www.broadinstitute.org/mitocarta/) and separated into nuclear-encoded genes and the 13 mtDNA-encoded protein-coding genes. Hallmark MYC Targets V1 and MYC Targets V2 were obtained from MSigDB using msigdbr. MitoCarta genes were excluded from the MYC signatures to avoid direct overlap with mitochondrial outcomes, and the remaining V1 and V2 genes were combined as a unique union to define nonmitochondrial MYC transcriptional activity. Paired tumor-normal differences in mitochondrial and MYC scores were evaluated using Wilcoxon signed-rank tests. In tumor-only analyses, nuclear mitochondrial and mtDNA scores were analyzed separately, with mitochondrial imbalance defined as the mtDNA score minus the nuclear mitochondrial score. Associations with MYC activity were assessed by Spearman rank correlation, with Pearson correlation and quartile-based comparisons used as complementary analyses.

For the MYC-ordered mitochondrial-state heatmap, predefined MSigDB gene sets were used: HALLMARK_OXIDATIVE_PHOSPHORYLATION; REACTOME_MITOCHONDRIAL_TRANSLATION, REACTOME_CITRIC_ACID_CYCLE_TCA_CYCLE, REACTOME_MITOCHONDRIAL_FATTY_ACID_BETA_OXIDATION, REACTOME_MITOCHONDRIAL_BIOGENESIS, and REACTOME_DETOXIFICATION_OF_REACTIVE_OXYGEN_SPECIES; and GOBP_MITOCHONDRIAL_RIBOSOME_ASSEMBLY and GOBP_ONE_CARBON_METABOLIC_PROCESS. The 13 mtDNA-encoded protein-coding genes were scored separately. Tumors were ordered by nonmitochondrial MYC activity without sample clustering.

Mitophagy and autophagy analyses used curated, non-overlapping gene modules representing receptor-mediated mitophagy, LC3/GABARAP ATG8 proteins, mitochondrial fission, core autophagy machinery, lysosome-autophagy output, and the PINK1/Parkin pathway. Modules were restricted to key pathway components to minimize redundancy among related transcriptional programs; exact gene memberships are provided in Supplementary Table 3. Multiple testing was controlled within analysis families using the Benjamini-Hochberg procedure. Data processing and visualization were performed in R 4.3 using tidyverse, ggplot2, ggrepel, and patchwork.

### Metabolomics

Flash frozen tissue was pulverized on liquid nitrogen (H37260-0100, Bel-Art Products). 40 vols of 40:40:20 (*v/v/v*, meOH: ACN: H2O) with 0.5% formic acid was added to samples, vortexed, placed on ice, and quenched with 8% ammonium bicarbonate. Protein was precipitated on dry ice, and samples centrifuged cold 3X at max speed. 100uL of sample was taken for LC-MS analysis. 200uL of sample was dried and used for GC-MS analysis.

For GC-MS analysis: dried metabolites were derivatized to form methoxime-TBDMS adducts by incubating with 1% methoxyamine hydrochloride (Sigma-Aldrich #226904) in pyridine at 70 °C for 15 minutes. Next, N-tert-Butyldimethylsiyl-N-methyltrifluoroacetamide (MTBSTFA, Sigma-Aldrich #375934) was added and incubated for 60 minutes. Derivatized samples were analyzed using an Agilent Technologies 8890 GC System with a DB-5MS GC column (Agilent #122-5532G) coupled to an Agilent Technologies 5977C mass spectrometer.

For LC-MS analysis: Samples were measured on the Q-Exactive Plus Orbitrap mass spectrometer coupled to a Vanquish Horizon Liquid Chromatography system (Thermo Scientific). A Waters XBridge BEH Amide Column (#186006724) with Waters BEH Amide XP VanGuard Cartridge (#186007763) was used. Mobile phase A was water:acetonitrile 95:5 (*v/v*) with 20mM ammonium acetate (Sigma-Aldrich #09689) + 20mM ammonium hydroxide (Sigma-Aldrich #09859), pH 9.45, with 5uM (NH4)2HPO4 (Thermo Fisher # AC201822500) and 50mM medronic acid (Millipore Sigma #1984-15-2). Mobile phase B was acetonitrile. Flow rate was 0.15ml/min throughout and the gradient over the course of 25 minutes went as follows: 0 min, 90%B; 2 mins, 90%B; 3 minutes, 75% B; 7 minutes, 75% B; 8 minutes, 70% B; 9 minutes, 70% B; 10 minutes, 50% B; 12 minutes, 50% B; 13 minutes, 25% B; 14 minutes, 25% B; 16 minutes, 0% B; 20.5 minutes, 0% B; 21 minutes, 90% B; 25 minutes, 90% B.

Injections were 5 µl in volume, and both negative and positive scans were performed over a mass range of 70 to 1000 m/z, with a resolution of 70,000. Automatic gain control was set at 3e6. Source ionization parameters were optimized with the spray voltage at 4.0 kV and in-source CID of 5.0eV, and other parameters were as follows: capillary temp at 425, S-lens level at 50, aux gas heater temperature at 400, Sheath gas at 35, aux gas at 10, and sweep gas flow at 2. Data was collected by the Thermo Fisher software Xcalibur (v4.5.474.0). Peak picking was performed in El-Maven for positive and negative ion mode, and clustering performed using Metaboanalyst (https://www.metaboanalyst.ca/).

### Proteomics Sample Preparation, Acquisition, and Analysis

Samples were lysed in 20 mM HEPES (pH 7.2) and sonicated twice for 8 s at 60% amplitude using a Branson saddle sonicator. Protein concentrations were measured using a reducing agent–compatible bicinchoninic acid assay (Pierce), and samples were normalized to equal protein concentrations. Proteins were reduced with 5 mM dithiothreitol (DTT) for 20 min at 60 °C. After cooling to room temperature, cysteines were alkylated with 20 mM *N*-ethylmaleimide (NEM) for 20 min at room temperature, and residual NEM was quenched with 10 mM DTT. Proteins were purified using a modified SP3 protocol (Hughes et al., 2019) with a 1:1 mixture of Sera-Mag SpeedBead carboxylate-modified E3 and E7 magnetic particles (Cytiva; catalog nos. 65152105050250 and 45152105050250). Samples were adjusted to 60% (v/v) ethanol and incubated for 1 h to bind proteins to the beads, after which the beads were washed four times with 80% (v/v) ethanol. Bead-bound proteins were digested overnight at 24 °C with LysC (20 ng µL⁻¹; Wako) in 2 M guanidine hydrochloride and 10 mM EPPS (pH 8.5) with agitation. Samples were then diluted fourfold with 10 mM EPPS and digested further overnight at 37 °C with LysC (20 ng µL⁻¹; Wako) and trypsin (10 ng µL⁻¹; Promega). Peptides were dried under vacuum, resuspended in 100 µL of 5% phosphoric acid, and desalted using Pierce C18 tips (Rappsilber et al., 2007).

Peptides were analyzed by direct injection using a Vanquish Neo UHPLC system coupled to an Orbitrap Astral mass spectrometer (Thermo Fisher Scientific). Peptides were separated at 350 nL min⁻¹ on an Aurora Ultimate C18 column (25 cm × 75 µm inner diameter, 1.7-µm particles, 120-Å pore size; IonOpticks) at 60 °C using water (A) and acetonitrile (B), each containing 0.1% formic acid and 2% DMSO. In the 70-min method, the proportion of B was increased from 2% to 4.8% over 5 min, to 26.4% by 55 min, and to 98% by 65 min. Ions were generated by positive-mode nano-electrospray at 2.0 kV with an ion-transfer-tube temperature of 290 °C. MS1 spectra were acquired in the Orbitrap at a resolution of 240,000 over *m/z* 380–980, with a normalized AGC target of 500% and a maximum injection time of 5 ms. DIA MS/MS spectra were acquired in the Astral analyzer using 300 nonoverlapping, placement-optimized 2-Th isolation windows spanning precursor *m/z* 380–980. Precursors were fragmented by higher-energy collisional dissociation at a normalized collision energy of 25, and fragment ions were recorded over *m/z* 150–2000 with a normalized AGC target of 500% and a maximum injection time of 3 ms.

Raw files were analyzed using DIA-NN v2.3.1 Academia (Demichev et al., 2020). A precomputed *in silico*–predicted spectral library derived from the *Mus musculus* and common-contaminant FASTA databases was used for the initial search. The first analysis pass generated a DIA-derived empirical library with empirically aligned retention times, after which all runs were reanalyzed against this library in a second pass with match-between-runs enabled. *In silico* digestion used Trypsin/P specificity—cleavage C-terminal to K or R, including before proline— with up to one missed cleavage. Peptides of 7–30 amino acids and precursor charge states of 1–4 were considered. The precursor and fragment ranges were restricted to *m/z* 380–980 and 200–1800, respectively. NEM modification of cysteine (+125.047679 Da; UniMod:108) was specified as fixed, whereas methionine oxidation (+15.994915 Da; UniMod:35) was specified as variable, with at most one variable modification per precursor. Peptidoform scoring and localization of methionine oxidation were enabled. Mass-accuracy and scan-window settings were determined automatically from the first run, and quantification was performed using QuantUMS in high-accuracy mode (Grossmann et al., 2026). Precursor identifications were filtered at a 1% false discovery rate, and protein groups were retained at a 1% global q-value; protein inference used proteotypic peptides only.

### NAD^+^/NADH Measurement

10 volumes of cold lysis buffer (1:1, 0.2N NaOH, 1% dodecyltrimethylammonium bromide: PBS) was added to each sample, vortexed, and centrifuged. NAD^+^ and NADH were then measured according to the manufacturer’s protocol (Promega # G9071). Briefly, NAD^+^ was isolated by heating at 60 °C in an acidic solution and NADH was isolated by heating at 75 °C.

### GSH/GSSG

10 mg of frozen tissue was homogenized in 10 volumes of saline and centrifuged at 4 °C for 10 minutes. 5 uL was mixed with 20 uL of glutathione lysis reagent and measured according to the manufacturer’s protocol (Promega #V6612).

### Enzymatic Activity Assays

OTC activity was measured as previously described.^48^ Briefly, tissue samples were homogenized in 10 volumes of saline and centrifuged at 4 °C. 10 ug of total protein was loaded into a 96 well plate. Ornithine (Sigma-Aldrich #O6503), triethanolamine (Sigma-Aldrich #90279), and carbamyl phosphate (Sigma-Aldrich #C4135) were added to each sample and incubated at 37 °C for 30 minutes. After incubation, the reaction was stopped by addition of phosphoric/sulfuric acid mixture (3:1) followed by addition of 2,3 butanedione monoxime (Acros Organics #150371000) and incubated at 95 °C for 30 minutes. The plate was then cooled to room temperature, centrifuged, and absorbance measured at 490 nm. Citrulline generation was fit to a standard curve and subsequent OTC activity determined.

Glutamate dehydrogenase (GDH) activity was measured according to the manufacturer’s protocol (ab102527). Briefly, tissue was homogenized in assay buffer and activity measured via a coupled NADH-linked assay. Glutamine synthetase (GLUL) activity was measured according to the manufacturer’s protocol (ab284572). Briefly, tissue was homogenized in assay buffer and activity measured via a coupled ATP-linked assay. Alkaline phosphatase (ALP) assay was measured according to the manufacturer’s protocol (ab83369). Briefly, tissue was homogenized in assay buffer and activity measured via dephosphorylation of p-nitrophenyl phosphate (pNPP).

### Mitochondrial Content

Mitochondrial content was determined by qPCR of genomic DNA using the following primers, as previously described.^49^

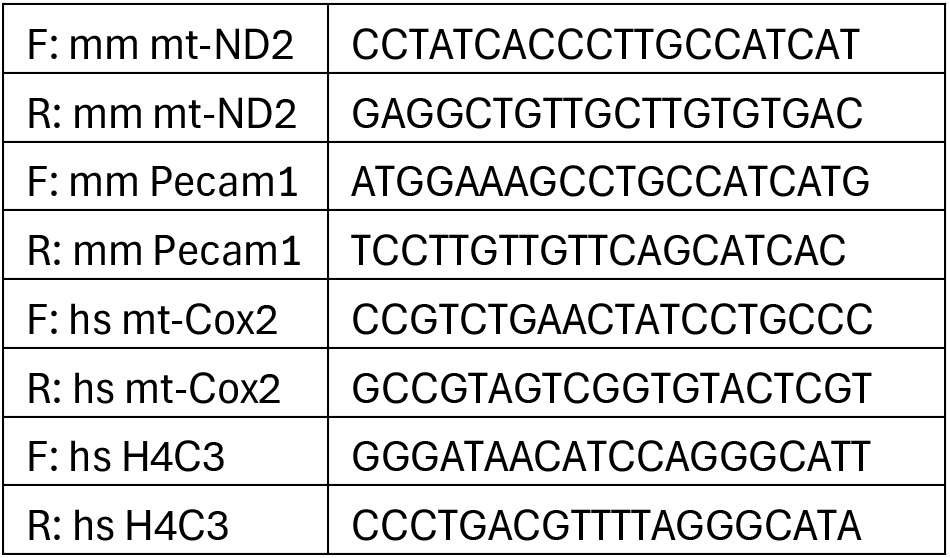

### Ammonia measurement

Plasma and intracellular ammonia was measured using a biochemical assay. Glutamate dehydrogenase (Sigma-Aldrich #G2626, 800U/mL) was incubated with 25 nmol alpha-ketoglutarate, 75 nmol NADH and incubated for 5 minutes. This solution was combined with 37.5 umol 2-oxopentanoic acid and 1.25 umol NADH and added to plasma or tissue homogenate. A plate reader was run in kinetic mode at 340nm for 30 minutes. Absorbance was converted to [NH3] using an ammonia standard curve.

### Cell Line Derivation

Cell lines were derived as previously described.^5^ Briefly, fresh tumors were minced and digested according to manufacturer’s protocol (Miltenyi Biotec #130-105-807). Cells were plated and passaged 8 times before use to ensure removal of non-tumor cells.

### Mitochondrial Respiration

Tissue was homogenized in mitochondrial isolation buffer [5mM HEPES, 70 mM sucrose, 220 mM mannitol, 5 mM KH2PO4, 5 mM MgCl2, 1 mM EGTA, pH 7.2] and mitochondria were isolated by differential centrifugation and examined for complex specific activity.^45,49^ 5ug of mitochondria were plated in a seahorse plate, allowed to settle for 10 minutes, then centrifuged at max speed to pellet onto the plate. Complex specific buffers were added and the seahorse immediately started. The mix:wait:measure cycle was (2:0:2 minutes) and injections as follow: port A: ADP (4mM), port B: oligomycin (2uM), port C: FCCP (2uM), port D: antimycin A (2uM) or azide (for complex IV, 50mM).

|  |  |
| --- | --- |
| Complex I | Pyruvate (10mM) / Malate (1mM) |
| Complex I | Glutamate (10mM) / Malate (1mM) |
| Complex II | Succinate (5mM) / Rotenone (0.002uM) |
| Complex IV | TMPD (0.1mM) / Ascorbate (10mM) / antimycin A (0.002mM) |

Homogenate respiration was measured from previously frozen tissue sections.^50^ Briefly, tumor sections were homogenized on ice using mitochondrial isolation buffer and centrifuged at 1,000 xg for 10 minutes at 4C and supernatant collected. Protein concentration was determined by BCA assay (Thermo Fisher). 10 ug of homogenate was loaded onto a Seahorse microplate in 20uL of isolation buffer and centrifuged at 2,000 xg for 2 minutes at 4C. Then 130 uL of isolation buffer containing cytochrome c [10 ug/mL] (Sigma-Aldrich #C2506) was added to each well and loaded into the Seahorse Analyzer. Substrate injection was as follows: NADH [1 mM], antimycin A [4 uM], TMPD/ ascorbate [0.5 mM and 1 mM, respectively]. Sodium azide [50 mM].

### Transmission Electron Microscopy (TEM)

Tissues for electron microscopic examination were fixed with 2.5% glutaraldehyde, 2.0% paraformaldehyde in 0.1M sodium cacodylate buffer, pH7.4, overnight at 4℃. After subsequent buffer washes, the samples were post-fixed in 2.0% osmium tetroxide with 1.5% K_3_Fe(CN)_6_ for 1 hour at room temperature and rinsed in dH2O prior to *en bloc* staining with 2% uranyl acetate. After dehydration through a graded ethanol series, the tissue was infiltrated and embedded in EMbed-812 (Electron Microscopy Sciences, Fort Washington, PA). Thin sections were stained with uranyl acetate and SATO lead and examined with a Talos L120C electron microscope fitted with a Ceta 16M digital camera.

## Data Availability

Proteomics data is included in supplemental table 1 and Data are available via ProteomeXchange with identifier PXD084050 (REVIEWER ACCESS: **Project accession:** and **Token:**). RNA-Sequencing data is included in supplemental table 2 (GSE148379). Further inquiries about resources and reagents should be directed to the lead contact, who will facilitate fulfillment.

## Author Contributions

**NPL:** conceptualization, data curation, formal analysis, validation, investigation, visualization, methodology, writing – original draft, writing – review and editing. **LCK:** investigation. **SDS:** data curation, investigation, visualization. **ML:** investigation. **XC:** investigation. **DZ:** investigation. **TP:** investigation. **CB:** investigation, resources. **BK:** writing – original draft, writing – review and editing. **MCS:** conceptualization, data curation, formal analysis, funding acquisition, visualization, methodology, supervision, resources, writing – original draft, writing – review and editing.

## Acknowledgements

The authors thank the Simon laboratory for comments and discussions on the manuscript. We thank the Institute of Structural Biology and Beckman Center for Cryo-EM at the University of Pennsylvania (RRID: SCR_022375), the Center for Molecular Studies in Digestive and Liver Diseases (P30DK050306, RRID: SCR_022420), and the Rodent Metabolic Phenotyping Core (S10-OD025098, RRID: SCR_022427). We thank Martin Wühr and Felix Keber for proteomics assistance. This work was supported by the Ludwig Cancer Research Institute (M.C.S), National Cancer Institute (NCI) R35CA220483 (M.C.S.), 5T32CA009140 (L.C.K. and N.P.L), Damon Runyon Postdoctoral Fellowship DRG2497 (N.P.L), and American Cancer Society Postdoctoral Fellowship PF-23-1034739-01-TBS (L.C.K). Biorender was used for the creation of some schematics.

## Supplemental Figure Legends

**Figure S1.** *MYC* Activity Correlates with Discordant Nuclear and mtDNA-Encoded Mitochondrial Gene Expression Across Cancers. **A)** Scatter plot of mean MitoCarta gene expression scores (as in Figure 1B) in matched primary tumors (y-axis) versus adjacent normal tissue (x-axis) across TCGA cancers (n = 684 total patients). Each point represents one patient, colored by cancer type; the dashed diagonal indicates equal tumor and normal expression (no difference). This panel is descriptive. Study codes are listed in Figure 1.

**B)** Bar chart of gene composition of Hallmark MYC Targets V1 and V2, showing retained non-mitochondrial genes (gray; n = 227) and excluded MitoCarta3.0-overlapping genes (red; n = 31). Excluded genes were removed to derive the non-mitochondrial MYC target score used throughout this study (Figures 1C–H).

**C)** Correlation between original (x-axis) and non-mitochondrial (y-axis) Hallmark MYC target scores in TCGA-LIHC (n = 371) confirming high concordance between the original and filtered scores. Spearman rho and unadjusted P values are indicated; the black line represents a linear fit.

**D)** Correlation of MYC mRNA expression (VST-normalized) and the non-mitochondrial MYC target score in TCGA-LIHC (n = 371), validating the score as a proxy for MYC transcriptional activity. Spearman rho and unadjusted P values are indicated; the black line represents a linear fit.

**E)** Scatter plot of mean non-mitochondrial MYC gene expression scores (as in Figure 1B) in matched primary tumors (y-axis) versus adjacent normal tissue (x-axis) across TCGA cancers (n = 684 total patients). Each point represents one patient, colored by cancer type; the dashed diagonal indicates equal tumor and normal expression (no difference). This panel is descriptive.

**F)** Paired nonmitochondrial MYC target scores in primary tumors and adjacent normal tissues across TCGA cancers. Lines connect matched samples, with black lines indicating cohort means; n, median Δ, and paired Wilcoxon signed-rank P values are shown. Study codes are listed in Figure 1.

**G)** Spearman correlations between *MYC* mRNA expression and individual mitophagy gene (*PGAM5*, *FUNDC1*, *NRF1*, *DNM1L* (DRP1); as identified in Figure 1H) expression in an independent cohort of HCC tissue (n = 15). Expression was measured by qRT-PCR and normalized to actin. Line represents a linear fit. Spearman rho and two-tailed p-values are shown.

**Figure S2.** MYC and Mitophagy Pathway Expression Predict Poor Survival in TCGA-LIHC. Kaplan-Meier overall survival curves for TCGA-LIHC patients (n = 371) stratified by high (Q4, top quartile) versus low (Q1, bottom quartile) expression; intermediate quartiles (Q2–Q3) were excluded. Statistical significance was assessed by log-rank (Mantel-Cox) test; P values are displayed in each panel. High-expression groups are shown in blue; low-expression groups are shown in red. Pathway-level analyses **(A–F)**: Pathway scores were calculated as ssGSEA

**A)** MYC Targets V1 (MSigDB Hallmark).

**B)** MYC Targets V2 (MSigDB Hallmark).

**C)** OXPHOS (MSigDB Hallmark).

**D)** Mitophagy (C2CP – Reactome Mitophagy).

**E)** PINK1-PARKIN-mediated mitophagy (C2CP – Reactome PINK1 PRKN Mediated Mitophagy).

**F)** Receptor-mediated mitophagy (C2CP – Reactome Receptor-Mediated Mitophagy). Gene-level analyses **(G–J)**: Individual gene mRNA expression from TCGA-LIHC RNA-seq data (VST-normalized). Genes were identified from mitophagy correlation analysis (Figure 1H).

**G)** *FUNDC1*.

**H)** *PGAM5*.

**I)** *NRF1*.

**J)** *DNM1L* (DRP1).

**Figure S3.** Loss of OXPHOS is a Conserved Feature of MYC-Driven HCC Across Strains and Genotypes. **A)** Volcano plot of VST-normalized RNA-Sequencing (GSE148379)^5^ in *MYC^OE^; PTEN^KO^* tumors compared to *CTNNB1^OE^; PTEN^KO^* tumors. Log_2_(Fold change) is plotted against the – log_10_(adjusted p-value) for each transcript. Significantly changing genes (log_2_(Fold change)>1 or < –1; Adjusted p-val <0.05) are colored black, mitochondrial genes (Mitocarta) are colored red, and nonsignificant genes are colored gray (n=3 per group).

**B)** Gene Set Enrichment Analysis (GSEA) running enrichment plot for the Hallmark OXPHOS gene set (MSigDB) from the RNA-Seq data shown in Panel A (*MYC^OE^; PTEN^KO^* vs. *CTNNB1^OE^; PTEN^KO^).* The normalized enrichment score (NES), nominal P value, and FDR q-value are displayed in the figure.

**C)** PCA clustering of VST-normalized RNA-Sequencing data (Panel A) of all indicated genotypes (n=3 per group). Genotypes and colors were as follows: Empty (black), *MYC^OE^; PTEN^KO^* (pink)*, MYC^OE^; CTNNB1^OE^* (green); *CTNNB1^OE^; PTEN^KO^* (purple). Variance for each principal component is shown in the figure.

**D)** PCA clustering of label-free proteomics (Figure 2E) of all indicated genotypes (n=3 per group). Genotypes and colors were as follows: Empty (black), *MYC^OE^; PTEN^KO^* (pink)*, MYC^OE^; CTNNB1^OE^* (green); *CTNNB1^OE^; PTEN^KO^* (purple). Variance for each principal component is shown in the figure.

**E)** PCA clustering of LC-MS (ESI+ and ESI-) metabolomics of all indicated genotypes (n=3 per group). Genotypes and colors were as follows: Empty (black), *MYC^OE^; PTEN^KO^* (pink)*, MYC^OE^; CTNNB1^OE^* (green); *CTNNB1^OE^; PTEN^KO^* (purple). Variance for each principal component is shown in the figure.

**F)** Representative western blot analysis of OXPHOS components (cV – ATP5A; cIII – UQCRC2; cIV – mtCO1; cII – SDHB; cI – NDUFB8) of the indicated genotype (Empty, *MYC^OE^; PTEN^KO^, MYC^OE^(no secondary driver))*. HSP60 (mitochondrial matrix), actin (whole cell lysate), and VDAC (outer mitochondrial membrane) were used as a loading control in MYC expressing tumors independent of a secondary driver (n=2 biological replicates per group). This panel is representative and not quantified.

**G)** Representative western blot analysis of OXPHOS components (cV – ATP5A; cIII – UQCRC2; cIV – mtCO1; cII – SDHB; cI – NDUFB8) of the indicated genotype (Empty, *MYC^OE^; TP53^KO^, MYC^OE^; PTEN^KO^, MYC^OE^; CTNNB1^OE^*, *CTNNB1^OE^; PTEN^KO^*, *CTNNB1^OE^; C-MET^OE^*) Actin was used as the loading control across MYC and β-catenin containing tumors (n= 2 per group). This panel is representative and not quantified.

**H)** Representative western blot analysis of OXPHOS components (cV – ATP5A; cIII – UQCRC2; cIV – mtCO1; cII – SDHB; cI – NDUFB8) of the indicated genotype (Empty, *MYC^OE^; TP53^KO^, MYC^OE^; PTEN^KO^)* in C57BL/6 and FVB/NJ mice. HSP90 was used as the loading control (n= 2 per group). This panel is representative and not quantified.

**Figure S4.** Impaired Urea Cycle Activity and Ammonia Accumulation Exacerbate Mitochondrial Dysfunction in HCC. **A)** Heatmap of log_2_ RNA expression (normalized to actin) of urea cycle and ammonia clearance genes (*CPS1, OTC, ASS1, ASL, ARG1, ARG2, NAGS, SLC7A1, GDH,* and *GLUL*) across genotypes as measured by qRT-PCR (n = 4-7 per group). Genotypes were as follows: Empty, *MYC^OE^; PTEN^KO^, MYC^OE^; CTNNB1^OE^*; *CTNNB1^OE^; PTEN^KO^*. Relative to empty vector, upregulated genes are colored red and downregulated genes are blue.

**B)** Representative western blot of urea cycle and ammonia clearance proteins (CPS1, ASS1, ASL, OTC, ARG1, GDH, and GLUL) from panel A across genotypes (n=2-3 per group). Genotypes were as follows: Empty, *MYC^OE^; PTEN^KO^, MYC^OE^; CTNNB1^OE^*; *CTNNB1^OE^; PTEN^KO^*. Tubulin was used as a loading control. This blot is representative and not quantified.

**C)** Heatmap of log_2_ TIC-normalized LC-MS metabolite abundance of urea cycle intermediates and related metabolites (arginine, aspartate, citrulline, fumarate, glutamate, glutamine, argininosuccinate, N-acetyl glutamate, and ornithine) from Figure S3E (n=5-7 per group). Genotypes were as follows: Empty, *MYC^OE^; PTEN^KO^, MYC^OE^; CTNNB1^OE^*; *CTNNB1^OE^; PTEN^KO^*. Relative to empty vector, increased metabolites are colored red and decreased metabolites are blue.

**D)** Heatmap of z-scores from top 50 (one-way ANOVA) altered metabolites from TIC-normalized LC-MS analysis across genotypes (n=5-7 per group). Genotypes were as follows: Empty, *MYC^OE^; PTEN^KO^, MYC^OE^; CTNNB1^OE^*; *CTNNB1^OE^; PTEN^KO^*. Color scheme was as follows: red (increased z-score) and blue (decreased z-score).

**E)** *Ex-vivo* glutamine synthetase (GLUL) activity (nmol/min/ ug protein) from snap frozen tumor tissue (n= 4-5 per group). Genotypes were as follows: Empty, *MYC^OE^; PTEN^KO^, MYC^OE^; CTNNB1^OE^*; *CTNNB1^OE^; PTEN^KO^*. Statistical significance was assessed by one-way Welch’s ANOVA with Dunnett’s T3 post-hoc test for multiple comparisons against the empty vector control.

**F)** *Ex-vivo* glutamate dehydrogenase (GDH) activity (nmol/min/ ug protein) from snap frozen tumor tissue (n= 4-5 per group). Genotypes were as follows: Empty, *MYC^OE^; PTEN^KO^, MYC^OE^; CTNNB1^OE^*; *CTNNB1^OE^; PTEN^KO^*. Statistical significance was assessed by one-way Welch’s ANOVA with Dunnett’s T3 post-hoc test for multiple comparisons against the empty vector control.

**G)** Relative abundance of ammonia transporter (RHBG) from proteomics Figure 2E (n=4-6 per group). Genotypes were as follows: Empty, *MYC^OE^; PTEN^KO^, MYC^OE^; CTNNB1^OE^*; *CTNNB1^OE^; PTEN^KO^*. Statistical significance was assessed by one-way Welch’s ANOVA with Dunnett’s T3 post-hoc test for multiple comparisons against the empty vector control.

**H)** Isolated liver mitochondria were treated with increasing NH_4_Cl concentrations (0, 0.2, 2 mM) for 10 minutes before being given succinate (cII substrate). Effect of ammonia concentrations on ADP-independent respiration (State IV OCR, left) and mitochondrial uncoupling (Respiratory control ratio (State III / State IV)) are shown (n=8 per group). Statistical significance was assessed by one-way Welch’s ANOVA with Dunnett’s T3 post-hoc test for multiple comparisons against the no ammonia control.

**I)** Isolated liver mitochondria were treated with increasing NH_4_Cl (0.2 or 2 mM) or NaCl (0.2 or 2 mM) concentrations for 10 minutes before being given pyruvate/malate (cI substrate). Respiratory control ratios (State 3_u_/State 4_o_) are shown (n=4-8 per group). Statistical significance was assessed by one-way Welch’s ANOVA with Dunnett’s T3 post-hoc test for multiple comparisons against the 0.2 mM NaCl control.

**J)** Mice were injected IP with NH_4_Cl (5mmol/kg) and sacrificed after 5 minutes. Liver mitochondria were isolated and function tested via Seahorse with succinate (complex II substrate). Mitochondrial uncoupling via the respiratory control ratio (State 3_u_/State 4_o_) is shown (n=6-8 per group). Statistical significance was assessed by Welch’s two-tailed t test.

**K)** Mitochondrial-OCR (basal – non mitochondrial, pmol/min/10k cells) in established tumor-derived cell lines of the indicated genotype (*MYC^OE^; PTEN^KO^* and *CTNNB1^OE^; PTEN^KO^*) treated with NH_4_Cl (0.25 mM) for 16 hours prior to Seahorse analysis. (n=4-8 per group). Statistical significance was assessed by Welch’s two-tailed t test.

**Figure S5.** Circulating Ammonia Alone is Insufficient to Recapitulate MYC-Driven Mitochondrial Phenotypes. **A)** Circulating ammonia levels (uM) in *MYC^OE^; PTEN^KO^* mice on a normal chow (20% protein, NC) and low protein (6% protein, LP) diet at endpoint (n=3-5 per group). Statistical significance was assessed by Welch’s two-tailed t test.

**B)** Kaplan-Meier survival curve with log-rank (Mantel-Cox) test of *MYC^OE^; PTEN^KO^* mice fed either a NC or LP diet (n=6 per group). P-values are displayed in the figure. Endpoint was defined when mice showed signs of morbidity and/or a >15% weight gain.

**C)** Representative western blot analysis of OXPHOS components (cV – ATP5A; cIII – UQCRC2; cIV – mtCO1; cII – SDHB; cI – NDUFB8) of *MYC^OE^; PTEN^KO^* mice fed NC or LP diets. Actin was used as the loading control (n= 4 per group). This panel is representative and not quantified.

**D)** % change from baseline body weight at diet initiation of non-tumor bearing mice on high ammonia diet (25% ammonium acetate *w/w*) compared to NC over 2 weeks (n=5 per group).

**E)** Liver to body weight ratios (%) in non-tumor bearing mice after 2 weeks of high ammonia or NC diet (n=5 per group). Statistical significance was assessed by Welch’s two-tailed t test.

**F)** Circulating ammonia (uM) and blood urea nitrogen (BUN, mg/dL) measured enzymatically in non-tumor bearing mice on high ammonia or NC diets (n=3-4 per group). Statistical significance was assessed by Welch’s two-tailed t test.

**G)** *Ex-vivo* ornithine transcarbamylase (OTC) activity (nmol/min/mg protein) in snap-frozen liver of non-tumor bearing mice on NC or high ammonia diets (n=5 per group). Statistical significance was assessed by Welch’s two-tailed t test.

**H)** TIC-normalized GC-MS TCA cycle metabolite (citrate, succinate, fumarate, malate, and aspartate) levels in non-tumor bearing, high ammonia diet liver relative to NC diet (n=5 per group). Statistical significance was assessed by Welch’s two-tailed t test.

**I)** Representative western blot analysis of OXPHOS components (cV – ATP5A; cIII – UQCRC2; cIV – mtCO1; cII – SDHB; cI – NDUFB8) and DRP1 of non-tumor bearing mice fed NC or high ammonia diets. Histone H3 and HSP60 were used as loading controls (n= 5 per group). This panel is representative and not quantified.

**Figure S6.** MYC Rather Than Tumor Differentiation Status Drives Mitochondrial Dysfunction in HCC. **A)** Representative (n=5 per group) H&E histology of HCC tumors showing well differentiated MYC tumors (*MYC^OE^; AXIN1^KO^).* Scale bar = 100 um.

**B)** *Ex-vivo* ornithine transcarbamylase (OTC) activity (nmol/min/mg protein) in snap-frozen liver of *MYC^OE^; AXIN1^KO^* tumors compared to empty vector controls (n=4-5 per group). Statistical significance was assessed by Welch’s two-tailed t test.

**C)** Circulating ammonia (uM) measured enzymatically in *MYC^OE^; AXIN1^KO^* tumor-bearing mice compared to empty vector controls (n=4-6 per group). Statistical significance was assessed by Welch’s two-tailed t test.

**D)** Representative western blot analysis of OXPHOS components (cV – ATP5A; cIII – UQCRC2; cIV – mtCO1; cII – SDHB; cI – NDUFB8) and DRP1 of *MYC^OE^; AXIN1^KO^* tumor-bearing mice compared to empty vector controls. Histone H3 and HSP60 were used as loading controls (n= 4-5 per group). This panel is representative and not quantified.

**E)** Seahorse analysis of complex II (succinate) and complex IV (TMPD/Ascorbate)-dependent respiration (OCR, pmol/min/μg protein) from frozen *MYC^OE^; AXIN1^KO^* tumor and adjacent normal tissue homogenates (n = 4–5 per group). Statistical significance was assessed by Welch’s two-tailed t test.

**F)** TIC-normalized GC-MS TCA cycle metabolite (citrate, succinate, fumarate, malate, and aspartate) levels in *MYC^OE^; AXIN1^KO^* tumors compared to empty vector controls (n=4-5 per group). Statistical significance was assessed by Welch’s two-tailed t test.

**Figure S7.** DRP1 Inhibition and Mitophagy Disruption Alter Mitochondrial Stress and Tumor Growth in MYC-Driven HCC **A)** Representative western blot of cytosolic cytochrome c in mitophagy knockout tumors following differential centrifugation (n = 2-3 per group). GAPDH was used as a cytosolic loading control. Genotypes were as follows: *MYC^OE^; PTEN^KO^* (sgCtrl*), MYC^OE^; PTEN^KO^; NRF1^KO^*(sgNRF1), *MYC^OE^; PTEN^KO^; FUNDC1^KO^* (sgFUNDC1*), MYC^OE^; PTEN^KO^; PGAM5^KO^* (sgPGAM5). This blot is representative and not quantified.

**B)** % change from baseline body weight of *MYC^OE^; PTEN^KO^* mice treated with DRP1 inhibitor (TAT-P110, 5mg/kg/day) or a control peptide (TAT) starting at 2 weeks post oncogene delivery (n=9 per group).

**C)** Kaplan-Meier survival curve with log-rank (Mantel-Cox) test of *MYC^OE^; PTEN^KO^* mice treated with DRP1 inhibitor (TAT-P110, 5mg/kg/day) or a control peptide (TAT) starting at 2 weeks post oncogene delivery (n=9 per group). P-values are displayed in the figure. Endpoint was defined when mice showed signs of morbidity and/or a >15% weight gain.

**D)** Representative (n=3 per group) IHC of total DRP1 expression in *MYC^OE^; PTEN^KO^; DNM1L^WT^ (*sgCtrl) vs. *MYC^OE^; PTEN^KO^; DNM1L^KO^ (sg*DRP1) tumors. Scale bar = 100 um.

**E)** Enzymatic quantification of redox ratio (NAD^+^/NADH) in tumors (n=6 per group). Genotypes were as follows: *MYC^OE^; PTEN^KO^; DNM1L^WT^* (sgCtrl) vs. *MYC^OE^; PTEN^KO^; DNM1L^KO^* (sgDRP1). Statistical significance was assessed using two-tailed Welch’s t test.

**F)** Enzymatic quantification of lipid peroxidation product malondialdehyde (MDA, uM/mg tissue) in tumors as measured by TBARS (n=6 per group). Genotypes were as follows: *MYC^OE^; PTEN^KO^; DNM1L^WT^* (sgCtrl) vs. *MYC^OE^; PTEN^KO^; DNM1L^KO^* (sgDRP1). Statistical significance was assessed using two-tailed Welch’s t test.

