## Supplemental Figures for "MYC Suppresses Mitochondrial Function in Hepatocellular Carcinoma through NRF1-Driven Receptor-Mediated Mitophagy"

Figure S1

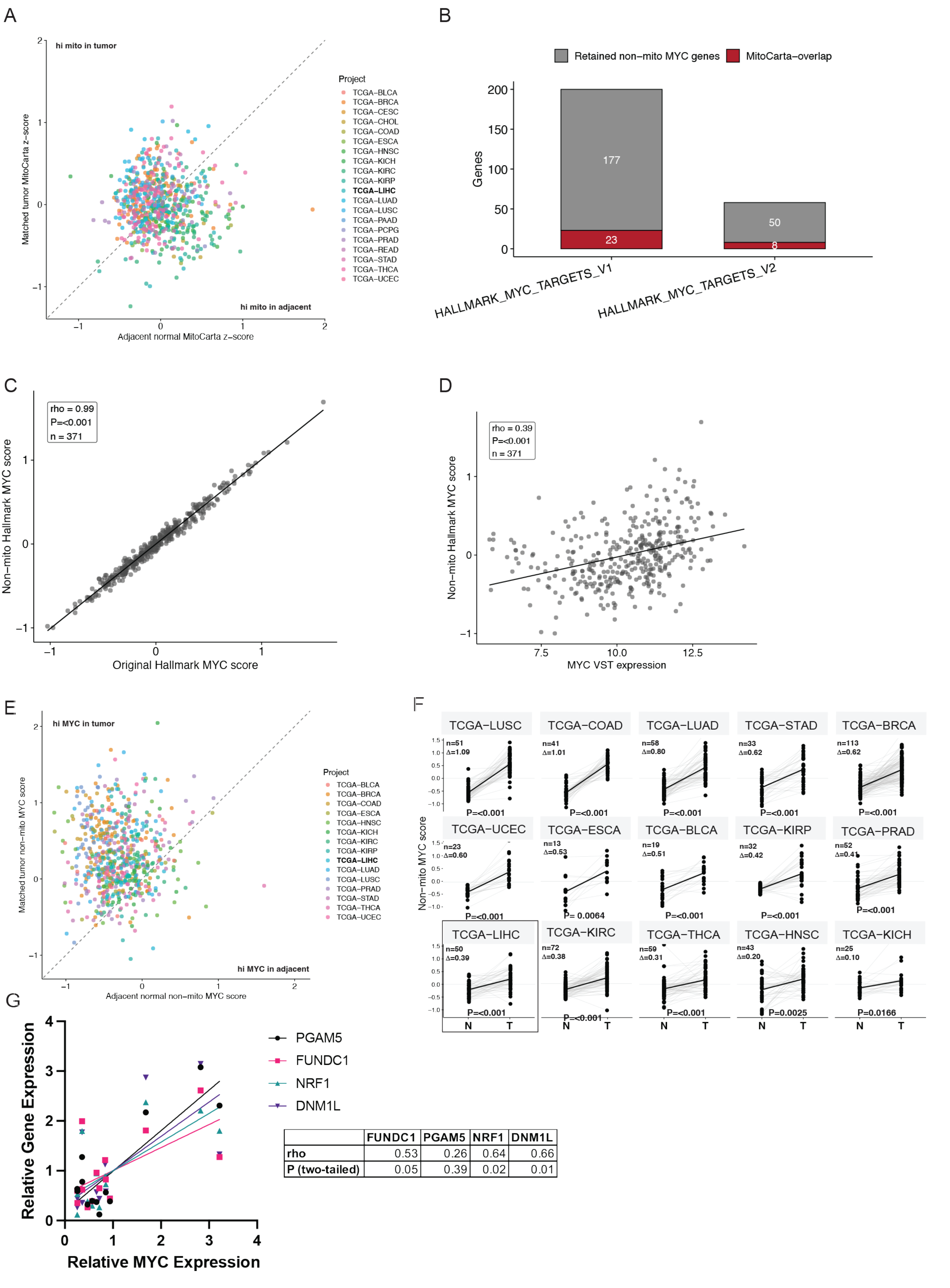


| | FUNDC1 | PGAM5 | NRF1 | DNM1L |
|---|---|---|---|---|
| rho | 0.53 | 0.26 | 0.64 | 0.66 |
| P (two-tailed) | 0.05 | 0.39 | 0.02 | 0.01 |

MYC Activity Correlates with Discordant Nuclear and mtDNA-Encoded Mitochondrial Gene Expression Across Cancers

Figure S2

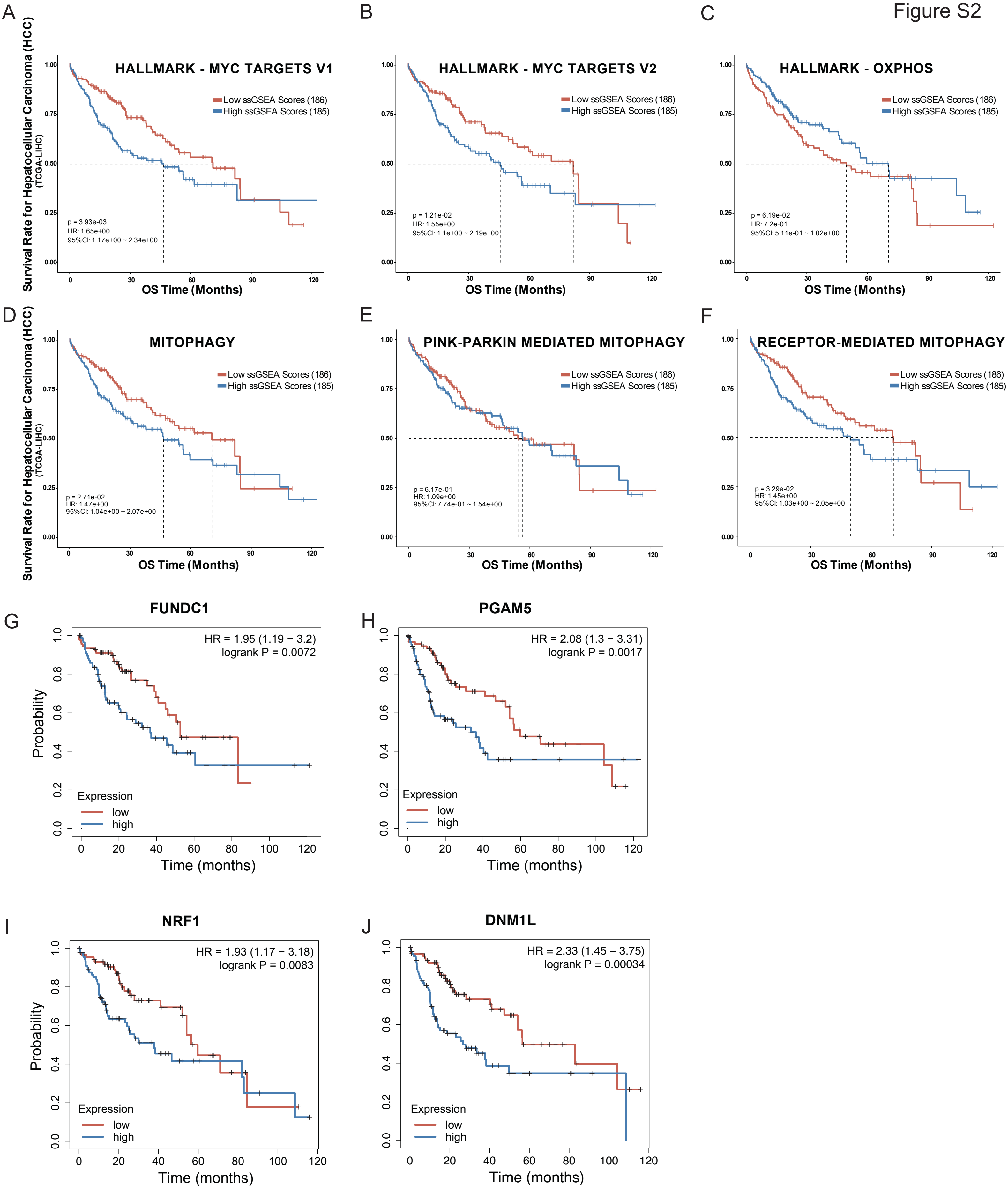


MYC and Mitophagy Pathway Expression Product Poor Survival in TCGA-LIHC

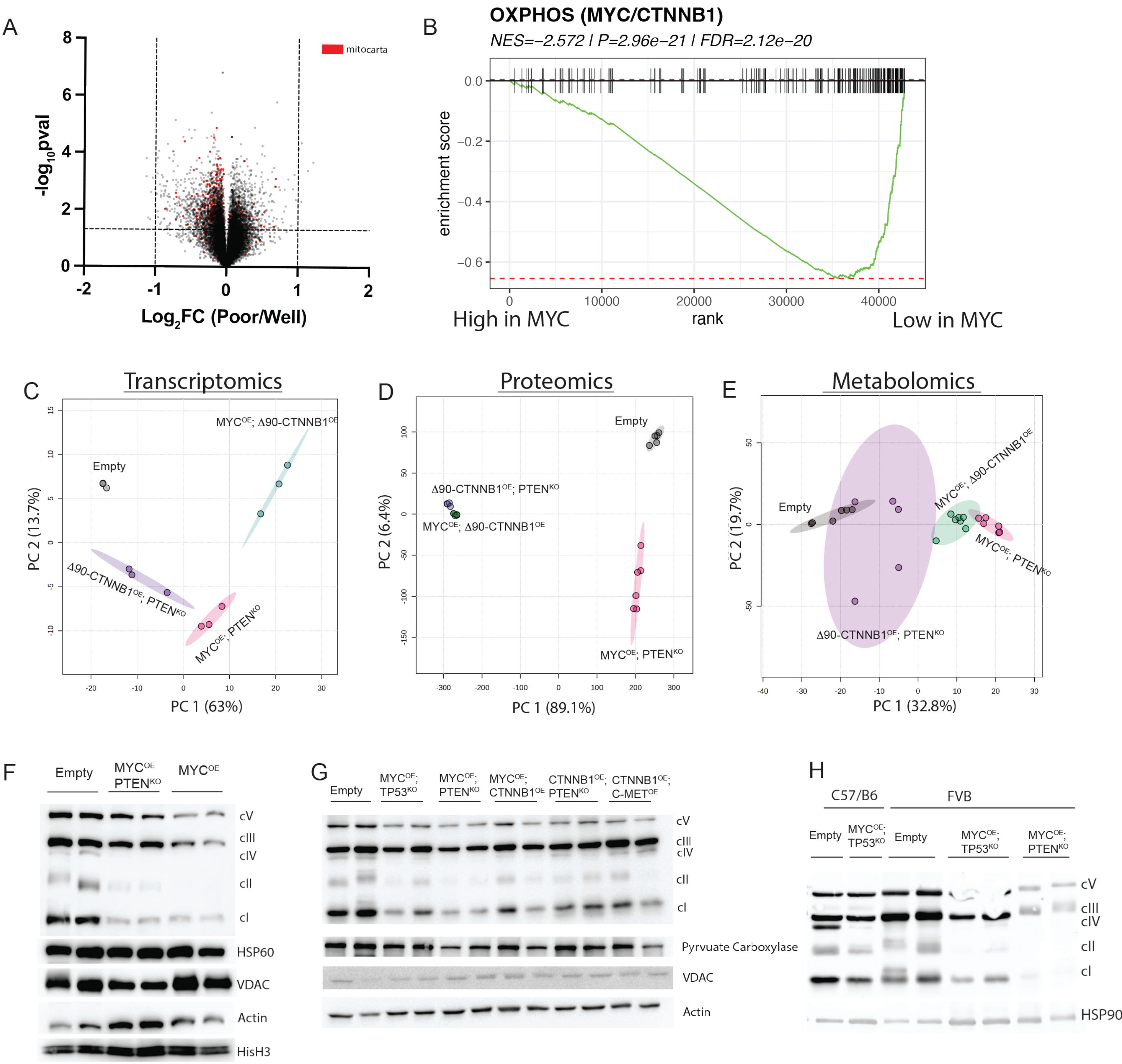


Loss of OXPHOS is a Conserved Feature of MYC-Driven HCC Across Strains and Genotypes

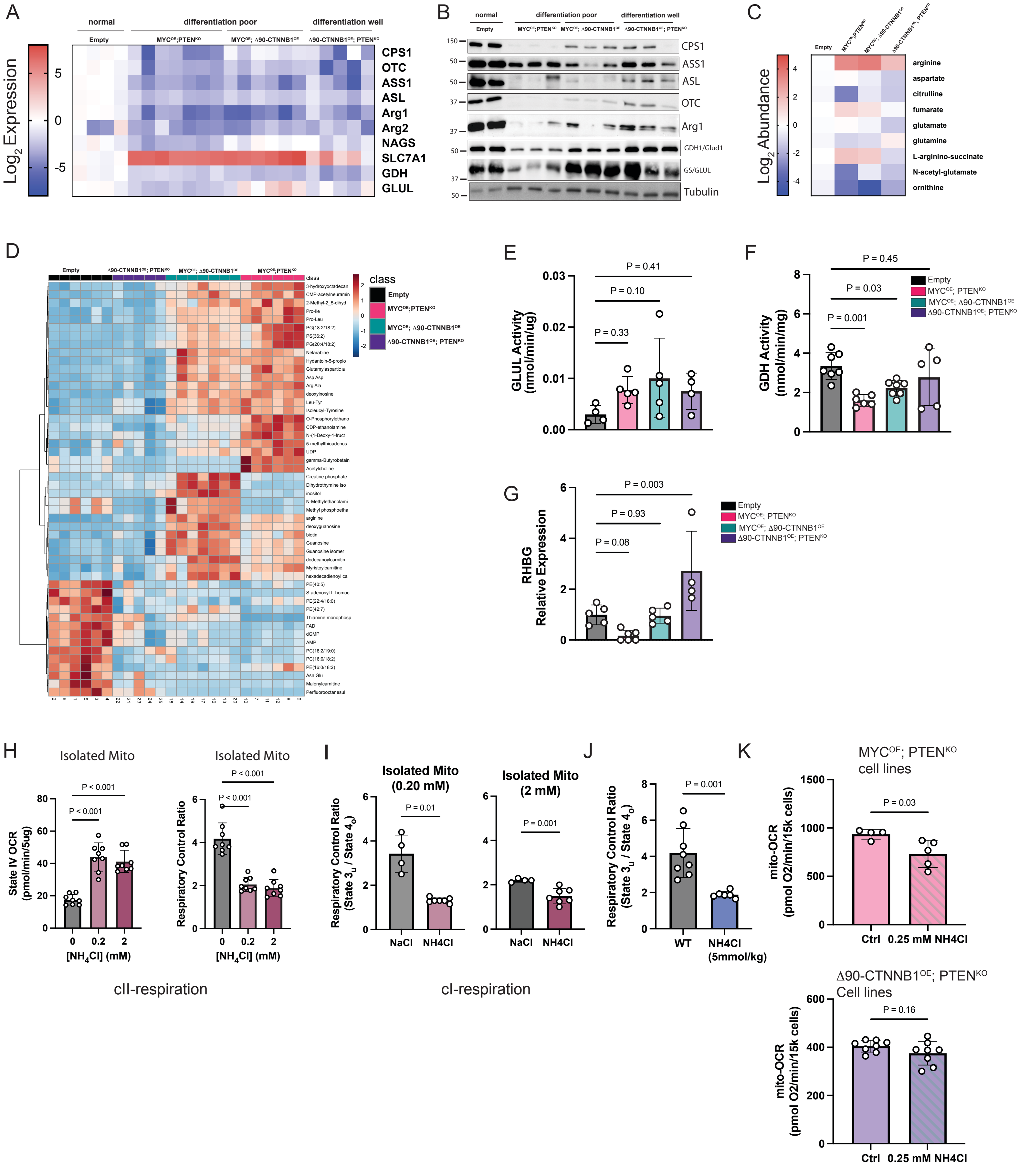


Impaired Urea Cycle Activity and AMmonia Accumulation Exacerbate Mitochondrial Dysfunction in HCC

Figure S5

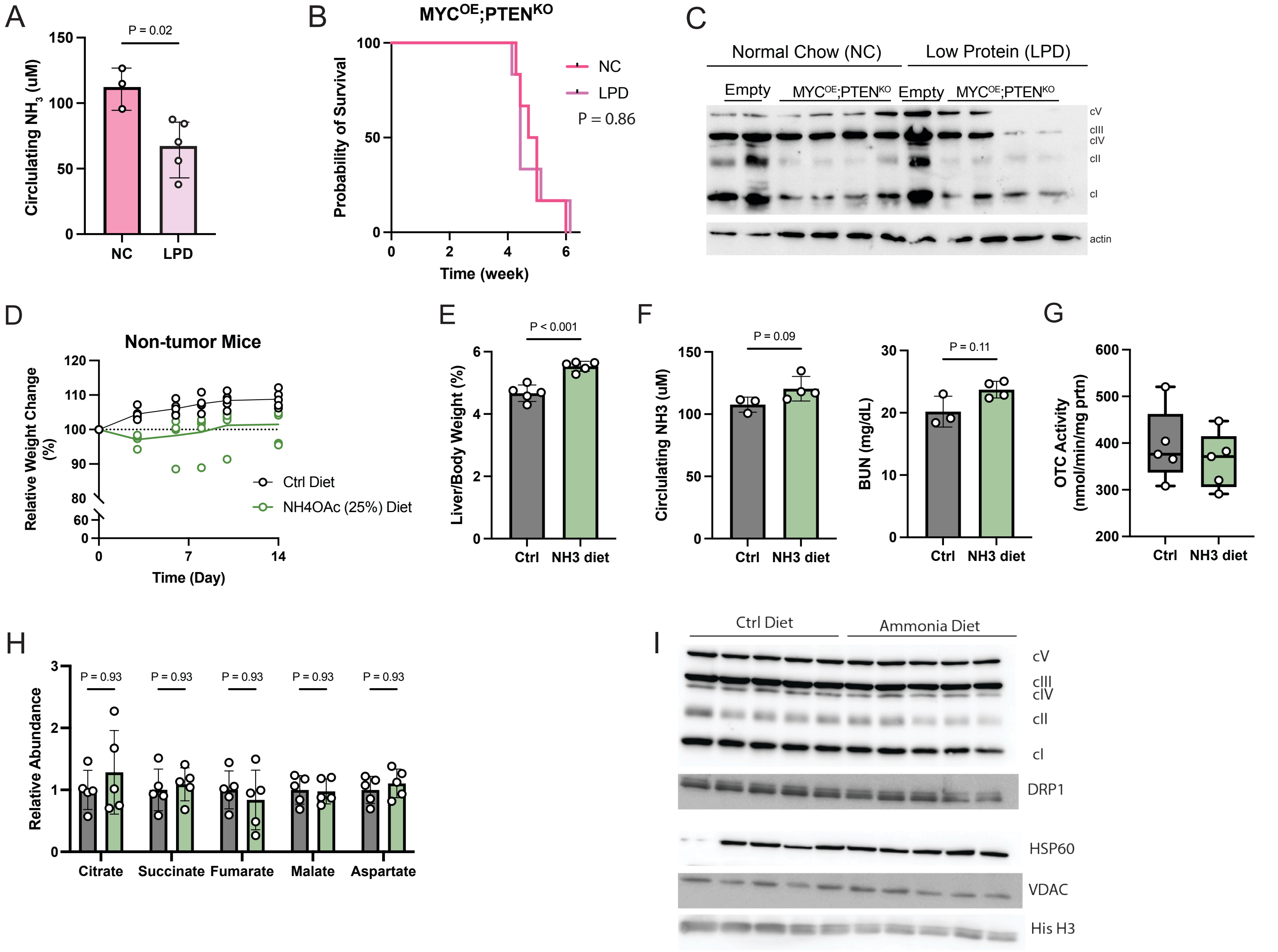


Circulating Ammonia Alone is Insufficient to Recapitulate MYC-Driven Mitochondrial Phenoytpes

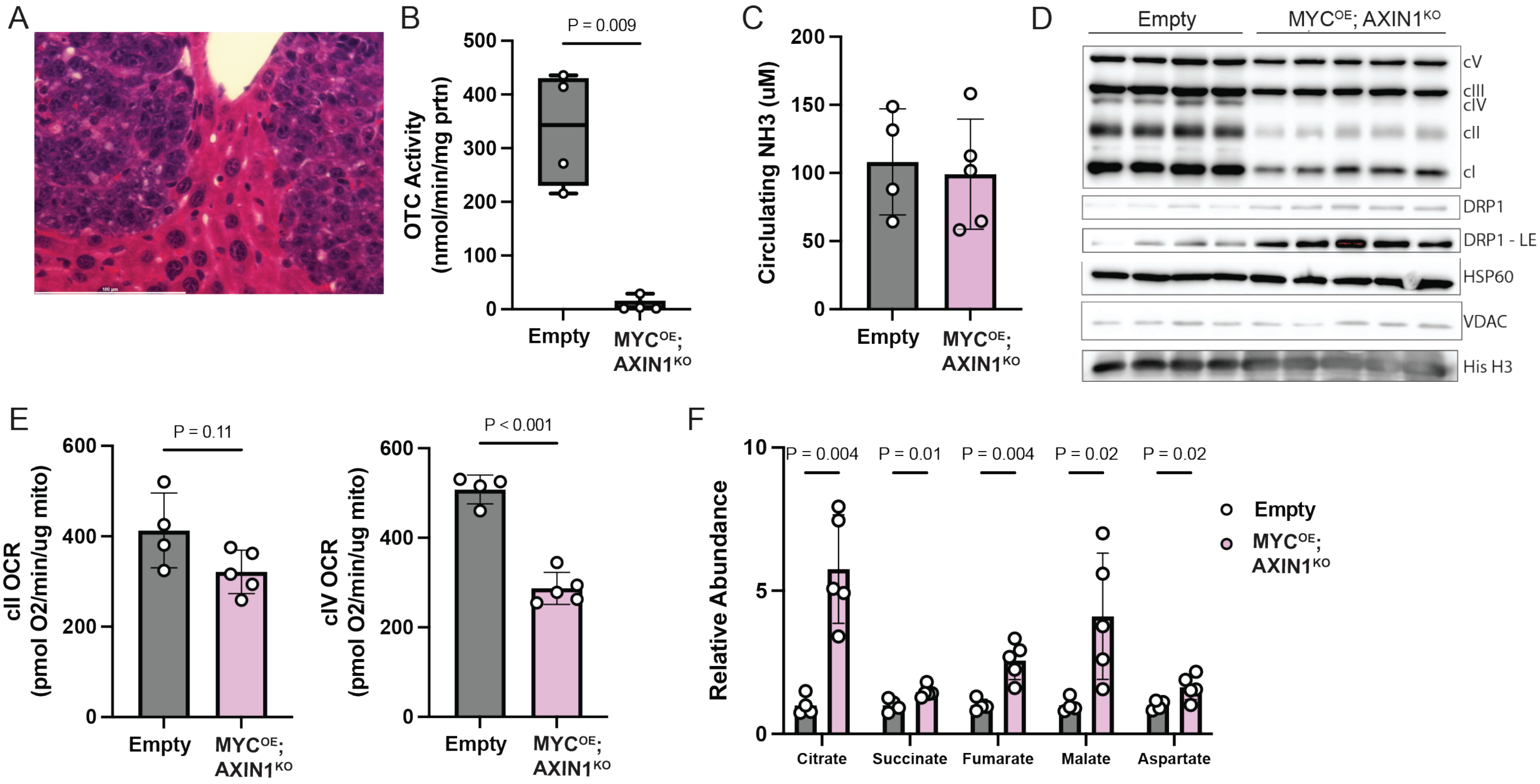


MYC Rather Than Tumor Differentiation Status Drives Mitochondrial Dysfunction in HCC

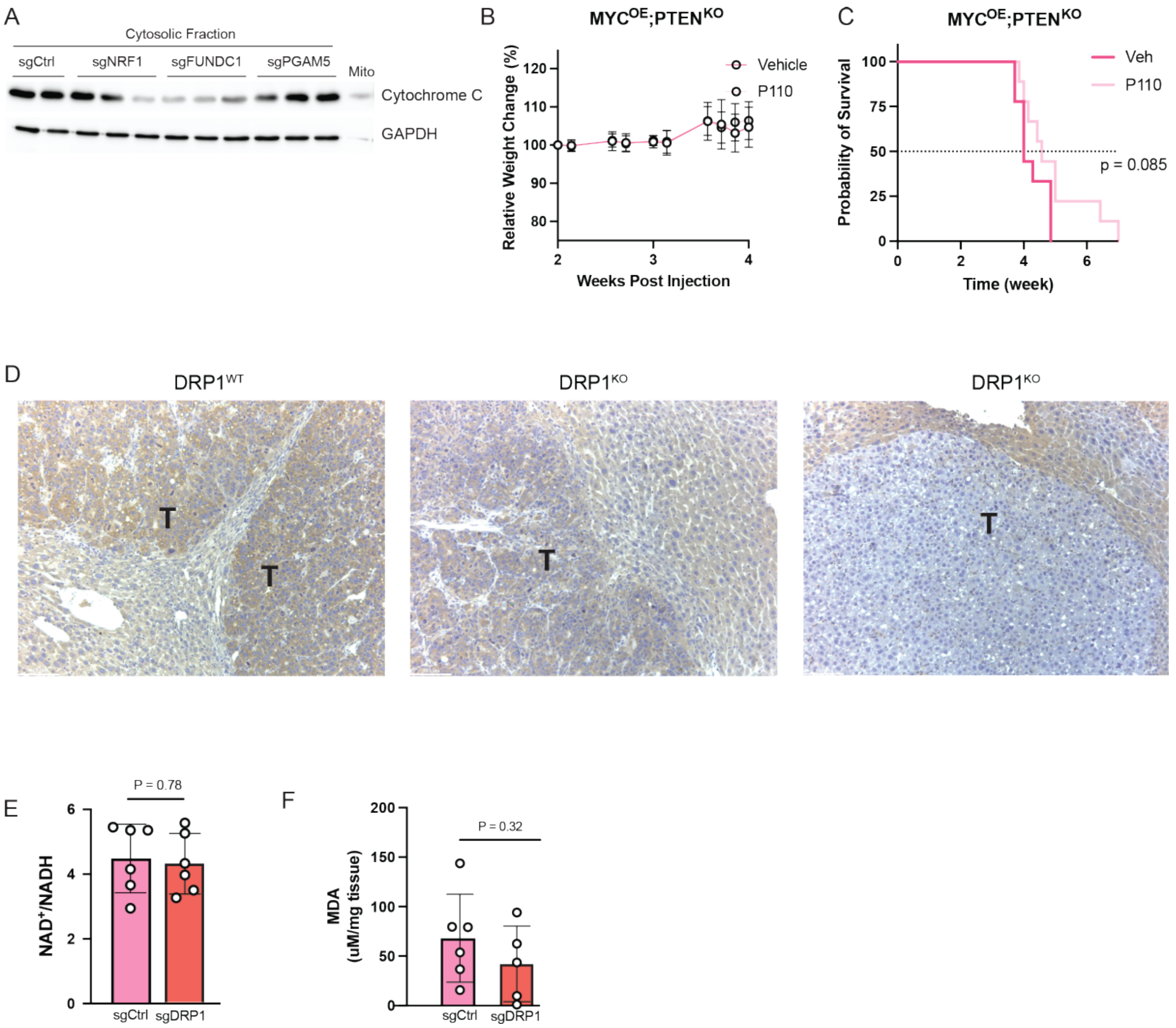


Pharmacologic Inhibition of DRP1 Suppresses Mitophagy and Reduces Tumor Burden in MYC-Driven HCC
